# The evolution of environmentally mediated social interactions and posthumous spite under isolation by distance

**DOI:** 10.1101/2023.09.22.558951

**Authors:** Charles Mullon, Jorge Peña, Laurent Lehmann

## Abstract

Many social interactions happen indirectly via modifications of the environment, e.g. through the secretion of functional compounds or the depletion of renewable resources. Here, we derive the selection gradient on a quantitative trait affecting dynamical environmental variables that feed back on reproduction and survival in a finite patch-structured population subject to isolation by distance. Our analysis shows that the selection gradient depends on how a focal individual influences the fitness of all future individuals in the population through modifications of the environmental variables they experience, weighted by the neutral relatedness between recipients and the focal. The evolutionarily relevant trait-driven environmental modifications are formalized as the extended phenotypic effects of an individual, quantifying how a trait change in an individual in the present affects the environmental variables in all patches at all future times. When the trait affects reproduction and survival through a payoff function, the selection gradient can be expressed in terms of extended phenotypic effects weighted by scaled-relatedness. We show how to compute extended phenotypic effects, relatedness, and scaled-relatedness using Fourier analysis, which allow us to investigate a broad class of environmentally mediated social interactions in a tractable way. We use our approach to study the evolution of a trait controlling the costly production of some lasting commons (e.g. a common-pool resource or a toxic compound) that can diffuse in space and persist in time. We show that indiscriminate posthumous spite readily evolves in this scenario. More generally whether selection favours environmentally mediated altruism or spite is determined by the spatial correlation between an individual’s lineage and the commons originating from its patch. The sign of this correlation depends on interactions between dispersal patterns and the commons’ renewal dynamics. More broadly, we suggest that selection can favour a wide range of social behaviours when these have carry-over effects in space and time.

## 1 Introduction

Organisms continually interact with one another in ways that significantly impact their survival and reproduction. Such social interactions are incredibly diverse. Still, they can usefully be classified as to whether they occur directly between individuals, such as grooming or fighting over resources, or as to whether they are indirectly mediated by environmental modifications, such as through the depletion or enrichment of resources, or the release or detoxification of pollutants [1]. Direct social interactions thus take place among contemporaries who are physically close to one another, while environmentally mediated interactions can extend much further in space and time. When environmental modifications have long-lasting and long-ranging effects, indirect social interactions may occur between individuals whose lifetimes show little or no overlap. This may lead to forms of trans-generational helping (e.g. when the underconsumption of a slowly renewable resource ensures healthy stock maintenance for future generations) or harming (e.g. when overconsumption leads to stock collapse and poor harvest in the future).

An extreme form of trans-generational social behaviour is posthumous altruism, which is a behaviour that results in a net reduction of the survival and/or reproduction of an actor to benefit only recipients living beyond the actor’s death. Several models have explored the biological scenarios that favour the evolution of posthumous altruism [2, 3]. But perhaps more striking is posthumous spite, which is a behaviour that results in a net reduction of the survival and/or reproduction of an actor to harm recipients that live only after the actor’s death. The conditions that may favour posthumous spite remain unclear. Understanding this, and more generally the evolution of environmentally mediated social behaviour, requires describing how selection shapes traits that underlie both direct and indirect social interactions.

The theory devoted to the evolution of quantitative traits influencing direct social interactions is well established (see, e.g. [4, 5, 6, 7, 8, 9]). One of the main contributions of this body of work has been to highlight the importance of limited dispersal for determining how selection shapes social traits in populations that are subdivided into finite groups and spatially structured [4, 5]. Under limited dispersal, stochastic demographic effects resulting from finite group (or patch or deme) size generate genetic associations, whereby individuals expressing the same traits may be more or less likely to interact directly with one another than with individuals expressing alternative traits. The importance of such associations is enshrined in the fact that the selection gradient on a quantitative trait can be expressed as a marginal (or gradient) form of Hamilton’s rule [4, 5]. This gradient, which captures the first-order effects of selection, is sufficient to determine the trait values towards which a population converges under mutation-limited evolution ([7], i.e. to characterise convergence stability [10]).

The marginal form of Hamilton’s rule is computationally attractive because all the necessary information about interactions is summarized in pairwise relatedness coefficients evaluated under neutrality (i.e. in the absence of selection or trait variation). This remarkable simplification makes the selection gradient tractable under realistic demographic assumptions, particularly in populations exhibiting isolation by distance (e.g. lattice models [11, 12, 5]). This has opened the door to understanding the evolution of multiple types of direct social interactions in such populations (e.g. helping and harming [13, 14, 15, 16, 17, 18, 19, 20]; sex ratio [21]; and dispersal [22, 23, 11, 24]).

In contrast, modelling the evolution of social interactions mediated by abiotic or biotic environmental variables is significantly more challenging in spatially structured populations. This is because computing the selection gradient in this case also requires computing the joint distribution of pairwise relatedness and environmental variables in the population (under neutrality [25, 5]). Generally, this distribution is the stationary solution to a high-dimensional stochastic dynamical system that is difficult to analyse or, in some cases, even to characterise. The challenge is apparent in models that allow for trait-driven changes in local demography. Even in the island model of dispersal, where the spatial structure is only implicit [26], there is typically no analytical solution to the distribution of demographic states within groups. As a result, evaluating the selection gradient on traits with effects on local demography and/or ecology is computationally challenging (e.g. [27, 28, 25, 29, 30]). This “curse of dimensionality” becomes even more acute under isolation by distance, as the size of the state space on which relevant environmental (or demographic) variables fluctuate blows up exponentially, with the selection gradient now requiring the distribution of states among as well as within groups (e.g. eq. 22 in [25]).

To circumvent this challenge, two approximations have been suggested. One is the pair approximation developed for lattice-structured populations, where typically at most one individual lives in sites connected by stepping-stone dispersal [31, 32, 33, 34, 35, 36, 37, 38]. Pair approximation is based on moment equations of the demographic state distribution that ignore third and higher-order moments. Under this approximation, the selection gradient can be written in the form of Hamilton’s marginal rule, thus allowing for a sharp understanding of some of the effects of demography on the evolution of social behaviour ([19, 20] see also [39]). However, pair approximation is not straightforward when considering arbitrarily complex dispersal patterns (e.g. [40]), patches with more than one individual, or trait-driven environmental state variables.

Another approximation relies on considering the dynamics of environmental state variables to be deterministic with a stable fixed point, so that there are no environmental stochastic fluctuations in the absence of genetic variation [41]. The selection gradient can then be readily expressed as a marginal Hamilton’s rule with inter-temporal fitness effects arising through trait-driven environmental modifications at different temporal distances. In addition to being simpler to compute than the original problem, this decomposition allows the delineation between a component of selection resulting from direct social interactions and a component arising indirectly through changes in the environmental dynamics. So far, this approach has been applied only to the island model—hence, in the absence of isolation by distance ([41, 42]. For populations exhibiting isolation by distance, there exist general, marginal Hamilton’s rule like, formulas. These express the selection gradient in terms of intertemporal and spatial effects of trait expression on the fitness of all possible recipients [2, 43]. However, how environmental modifications mediate the long-lasting and long-ranging fitness effects due to trait expression remains implicit in these general formulas. To better understand selection resulting from indirect social interactions via environmental feedback, fitness effects need to be unpacked in terms of trait-driven environmental modifications at different temporal and spatial distances.

Here, we do exactly that. We fully characterise the selection gradient on a trait that impacts the deterministic dynamics of environmental state variables that can be abiotic or biotic, and which feed back on the survival and reproduction of the evolving species in a general model of isolation by distance. Using Fourier analysis, we express this gradient in terms of extended phenotypic effects and relatedness coefficients scaled to local competition, both of which provide biological insights about the nature of selection and are straightforward to compute for a wide range of classical evolutionary models (e.g. the Wright-Fisher model and the Cannings model). We use our results to investigate the evolution of environmentally mediated helping and harming through space and time, such as via the production of a lasting common-pool resource or the release of a durable toxic compound. Our analyses indicate that indiscriminate spite where individuals suffer a cost to harm others living in the future, even beyond the actor’s death, readily evolves by natural selection. This result is in contrast with previous findings suggesting that indiscriminate spite rarely evolves when it occurs through direct interactions. More broadly, our model and results suggest that selection can favour a wide range of social behaviours when these are mediated in space and time through environmental feedbacks.

## 2 Models

### 2.1 Spatial structure, life cycle, traits and environmental variables

We consider a population of homogeneous individuals subdivided among *D* homogeneous patches (or demes or groups), each carrying *N* adult individuals (see Table 1 for a list of the key symbols used). The population is censused at discrete demographic time steps between which the following events occur in cyclic order: (a) reproduction and adult survival; (b) dispersal among patches; and (c) density-dependent regulation within patches such that each patch contains *N* adult individuals at the beginning of the next demographic time step.

**Table 1.** Key general symbols.

| Table 1. Key general symbols |  |
| --- | --- |
| $D$ | number of patches in the population (section 2.1) |
| $N$ | number of adult individuals per patch (section 2.1) |
| $G$ | set of patches in the population (section 2.1) |
| $z_{\bullet}$ | trait value of the focal individual, who lives in patch 0 and time $t = 0$ (section 2.2) |
| $z_{\mathbf{k},t}$ | average trait of individuals *other* than the focal in patch $\mathbf{k}$ at time $t$ (section 2.2) |
| $n_{\mathbf{k},t}$ | environmental state variable in patch $\mathbf{k}$ at time $t$ (section 2.2) |
| $g$ | transition map determining the dynamics of the environmental state variables (eq. 2) |
| $w$ | individual fitness function, i.e. expected number of successful offspring (including the surviving self) produced by an individual over one demographic time step (eq. 1) |
| $s(z)$ | selection gradient (section 4) |
| $s_w(z)$ | selection due to intra-temporal effects (section 4) |
| $s_e(z)$ | selection due to inter-temporal effects (section 4) |
| $R_{\mathbf{k},t}$ | relatedness coefficient between the focal individual and another individual from patch $\mathbf{k}, t$ (eq. 7) |
| $m_{\mathbf{k}}$ | probability that an individual disperses to a patch at distance $\mathbf{k}$ from its natal patch (section 2.1) |
| $\mathcal{M}(\mathbf{h})$ | Fourier transform (eq. I.B in Box 1) of $m_{\mathbf{k}}$ |
| $\bar{\chi}_{\mathbf{k}}(\mathbf{h})$ | conjugate of character function (eq. I.E in Box 1) |
| $\mathcal{L}_{\mathbf{k}}(\mathcal{F})$ | inverse transform of $\mathcal{F}$ at $\mathbf{k}$ (eq. I.D in Box 1) |
| $e_{\mathbf{k},t}$ | extended phenotypic effect of the focal individual on the environmental state variable in patch $\mathbf{k}$ at time $t$ in the future (eqs. 10 and 14) |
| $\psi_{\mathbf{k}}$ | focal individual's effect on the environmental state variable of patch $\mathbf{k}$ over one generation (eq. 12) |
| $\Psi(\mathbf{h})$ | Fourier transform (eq. I.B in Box 1) of $\psi_{\mathbf{k}}$ |
| $c_{\mathbf{k}}$ | effect of the environmental state variable of one patch on the environmental state variable of another patch at distance $\mathbf{k}$ over one generation (eq. 13) |
| $\mathcal{C}(\mathbf{h})$ | Fourier transform (eq. I.B in Box 1) of $c_{\mathbf{k}}$ |
| $\pi$ | individual payoff function (section 3.3.1) |
| $\lambda_{\mathbf{k}}$ | coefficient of fitness interdependence through payoffs (eq. 20) |
| $\kappa_{\mathbf{k},t}$ | scaled relatedness coefficient between the focal individual and another individual from patch $\mathbf{k}, t$ (eq. 19) |
| $p_{\mathbf{k},t}$ | probability that a gene descending from the focal individual is in patch $\mathbf{k}$ at $t > 0$ steps in the future (eq. 22) |
| Key symbols from example section 3.4 |  |
| $B, \alpha_B$ | parameters tuning the effects of the environmental variable on the payoff of the focal individual (eq. 25) |
| $C, \alpha_C$ | parameters tuning the effects of the trait $z_{\bullet}$ of the focal individual on the payoff of the focal individual (eq. 25) |
| $P$ | commons production function (eq. 26) |
| $\epsilon$ | commons decay rate (eq. 26) |
| $d_{\mathbf{k}}$ | probability that a unit of commons moves to a patch at a distance $\mathbf{k}$ from its source patch (eq. 26) |
| $\mathcal{D}(\mathbf{h})$ | Fourier transform (eq. I.B in Box 1) of $d_{\mathbf{k}}$ |
| $q_{\mathbf{k},t}$ | probability that a non-decaying unit of the commons modified in the focal patch is located in patch $\mathbf{k}$ $t$ generations in the future (eq. 29) |
| $\Omega$ | expected genetic value in units of payoffs of all the individuals in the future that are affected by a unit of the commons in the focal patch (eq. 31) |

Patches are arranged homogeneously in *d* dimensions, with *D*_*j*_ patches in dimension *j* ∈ {1, …, *d*}. For example, under a lattice structure in a one-dimensional habitat, *D* = *D*_1_ patches are arranged on a circle, while in a two-dimensional habitat, *D* = *D*_1_ × *D*_2_ patches are arranged on a torus. We denote by 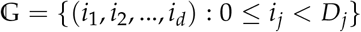 the set of all patches. The fact that patches are arranged homogeneously means that, at a technical level, we can endow the set 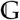 with an abelian group structure (see e.g. [44] for the use of such group structure in evolutionary biology), which will allow us to leverage techniques from Fourier analysis on finite abelian groups (Box 1).

#### Box 1

**Fourier analysis on finite abelian groups**

We assume that the set of patches 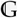 is endowed with an abelian group structure, which allows us to consider more general spatial structures than just lattice models (e.g. hierarchical structures). The group 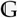 is defined as the direct product,

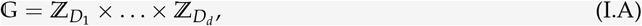

where 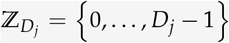 is the additive group of integers modulo *D*_*j*_. The group 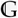 then consists of the set of all vectors **i** = (*i*_1_, …, *i*_*d*_) with 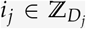 together with addition (where addition between two vectors is component-wise). On such a group, the Fourier transform ℱ(**h**) of function *f* at **h** ∈ 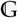 is given by

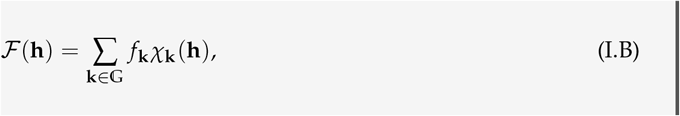

where the “character” function

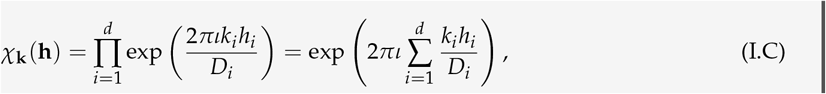

with 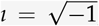, is defined for all 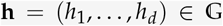 and 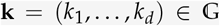. Here, we followed the convention of population genetics (e.g. [46, 47, 5]) and defined the Fourier transform in terms of the character *χ*_**k**_ (**h**) (instead of its conjugate, given in eq. I.E, which is more standard in mathematics and engineering). As such, the Fourier transform of *f* gives the characteristic function of *f* when *f* is a probability distribution. For instance, the Fourier transform 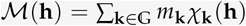 is the characteristic function of the dispersal distribution *m*_**k**_. The original function is recovered by using

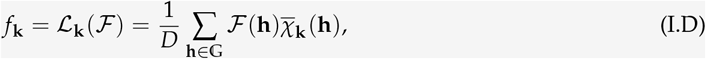

where ℒ_**k**_ (ℱ) is the inverse transform of ℱ at 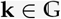, which is defined in terms of the conjugate of *χ*_**k**_ (**h**):

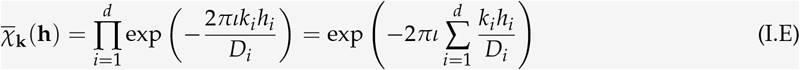

(e.g. [82]). Another property that we use in our analysis is the orthogonality relation between characters, i.e., the identity

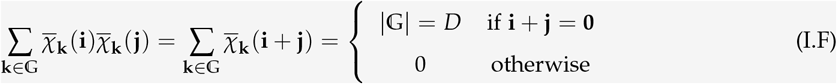

(see [82], p. 169).

Each patch is characterized by a quantitative state variable representing a biotic or abiotic environmental factor, which we refer to as an environmental state variable (e.g., density of a common-pool resource or pollutant, habitat quality). Meanwhile, each individual in the population is characterised by a genetically determined quantitative trait (e.g. consumption of a resource, release of a pollutant, investment into habitat maintenance) that influences the environment and the individual’s survival and reproduction. We are interested in the evolution of this trait under the following three assumptions.

1. *Trait and environmentally mediated reproduction and survival*. By expressing the evolving trait, individuals can directly affect the survival and/or reproduction of any other individual in the population. For example, individuals may engage in costly fights for resources in other patches and return to their own patch to share these resources with patch neighbours. The effects of trait expression on others are assumed to be: (a) spatially invariant, i.e. the marginal effect of an individual from patch 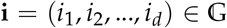 on the survival and/or reproduction of an individual in patch 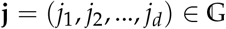 only depends on the “distance” **j** − **i** between the two patches (where **j** − **i** is calculated from the abelian group operation, see Box 1); and (b) spatially symmetric, i.e. the marginal effect of an individual from patch **i** on an individual in patch **j** is equal to the effect from **j** to **i**. We refer to these two characteristics (a) and (b) together as spatial homogeneity. The survival and reproduction of an individual may also depend on the environmental state variable of each patch, also in a spatially homogeneous way (i.e. the marginal effect of the environmental state variable of a patch **i** on the survival and reproduction of an individual residing in patch **j** only depends on the distance **j** − **i**, and is equal to the effect from **j** to **i**).
2. *Dispersal*. Each individual either stays in its natal patch or disperses to another patch. Dispersal occurs with non-zero probability so that patches are never completely isolated. We assume that dispersal is spatially homogeneous, i.e. that the probability of dispersal from one patch **i** to another **j** depends only on the distance **j** − **i** = **k** between the two patches (spatial invariance), and is equal to the probability of dispersing the distance **i** − **j** = − **k** (spatial symmetry). We can thus write *m*_**k**_ = *m*_−**k**_ for the probability that an individual disperses to a patch at a distance **k** from its natal patch (with 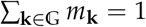).
3. *Trait and environmentally mediated environmental dynamics*. Through trait expression, individuals can affect environmental state variables from one demographic time step to the next. For example, the environmental variable may be a common-pool resource that individuals absorb locally, or a pollutant produced by individuals which then diffuses in the environment. Such trait effects on the environment are also spatially homogeneous (i.e. the marginal effect of an individual from patch **i** on the environmental state variable of patch **j** only depends on the distance **j** − **i** and is equal to the effect from **j** to **i**). These trait-driven environmental modifications can thus lead to inter-temporal, environmentally mediated social interactions.

### 2.2 The focal individual, its fitness, and environmental dynamics

The spatial homogeneity underlying all processes described above means that the patch indexed as 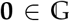 can be taken as a representative patch, and that any individual in this patch can be taken as a representative individual from the population. We refer to this patch and to this individual as, respectively, the focal patch and the focal individual. In general, we refer to patch 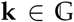 at time (or “generation”) *t* ≥ 0 prior to the focal generation as patch “**k**, *t*”. Thus, the focal patch corresponds to patch **0**, 0. In the following, we introduce some notation to describe trait and environmental variation in the population (see Fig 1C for a summary diagram of our model). We denote by *z*_•_ the realized value of the trait of the focal individual, and by *z*_**k**,*t*_ the realized average trait of individuals *other* than the focal individual living in patch **k**, *t* [e.g. for a one-dimensional habitat, *z*_1,1_ is the average trait expressed in the adjacent patch (patch 1) one time point before the focal generation]. Hence, *z*_**0**,0_ denotes the average phenotype among the patch neighbours of the focal individual in the focal generation (thus excluding the focal from the average). We use 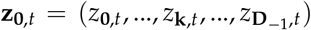 to denote the vector collecting all such realized phenotypes in 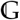 in lexicographic order, finishing with position **D**_−1_ = (*D*_1_ − 1, *D*_2_ − 1, …, *D*_*d*_ − 1). Finally, we use *n*_**k**,*t*_ to denote the environmental state variable in patch **k**, *t*; we collect the environmental state variables across all patches in the vector 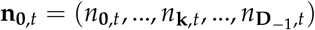.

**Figure 1:**
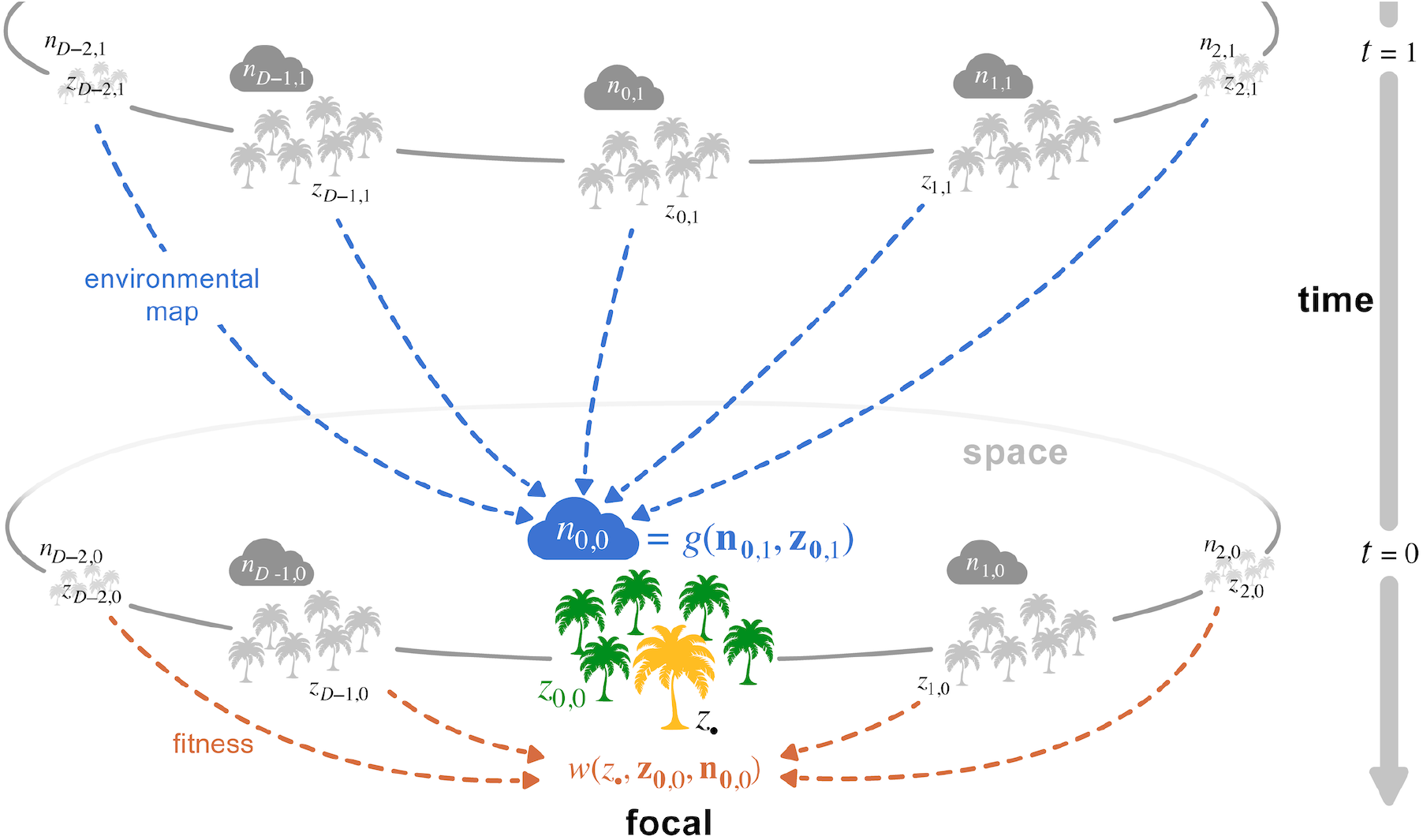
A model for environmentally mediated social interactions in space and time. Schematic description of our model for a one-dimensional lattice habitat (see sections 2.1-2.2 for details). Each patch *k* ∈ {…, *D* − 1, 0, 1, …} at time *t* ∈ {0, 1, …} in the past is characterized by an environmental state variable *n*_*k,t*_ (represented here by a cloud, e.g. water level, concentration of a pollutant, density of a resource), and the average trait value *z*_*k,t*_ expressed by the individuals it carries (e.g. water absorption rate, detoxifying capacity, handling time; individuals represented here as palms). The environmental state *n*_0,0_ of the focal patch *k* = 0 at time *t* = 0 depends on all environmental states and traits of the previous generation according to the environmental map *g* (blue dashed arrows, eq. 2). In turn, the fitness of a focal individual with trait *z*_•_ (in yellow) depends on all environmental states and traits expressed in its own generation according to the fitness function *w* (orange arrows, eq. 1). The two functions *g* and *w* thus characterise how evolutionary and environmental dynamics interact with one another through dual inheritance of traits and environmental state variables.

The fitness of the focal individual is determined by the function *w* : ℝ× ℝ^*D*^ × ℝ^*D*^ → ℝ_+_ such that

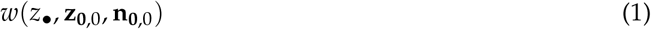

is the expected number of successful offspring (i.e. offspring that establish as adults, including the surviving self) produced over one demographic time by the focal individual with trait *z*_•_, when the trait average among other individuals at the different spatial positions is **z**_**0**,0_, and environmental state variables are **n**_**0**,0_. These state variables are obtained from the solution to the system of equations

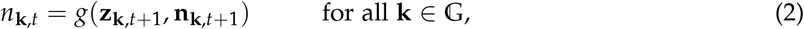

where *g* : ℝ^*D*^ × ℝ^*D*^ → ℝ is a transition map determining the dynamics of the environmental state variables **n**_**k**,*t*_ of all patches, which is a circular permutation of **n**_**0**,*t*_ with *n*_**k**,*t*_ as first element [e.g. for a one-dimensional lattice where *d* = 1, **n**_0,*t*_ = (*n*_0,*t*_, *n*_1,*t*_, …, *n*_*D* − 1,*t*_), **n**_1,*t*_ = (*n*_1,*t*_, *n*_2,*t*_, …, *n*_0,*t*_), **n**_2,*t*_ = (*n*_2,*t*_, *n*_3,*t*_, …, *n*_0,*t*_, *n*_1,*t*_), and so on]. The map *g* depends on (i) the traits in the whole population expressed at the previous generation via **z**_**k**,*t*+1_ (recall that *t* goes back in time), which is a circular permutation of the elements of **z**_**0**,*t*+1_ with *z*_**k**,*t*+1_ as first element [e.g. for a one dimensional lattice, **z**_0,*t*+1_ = (*z*_0,*t*+1_, *z*_1,*t*+1_, …, *z*_*D* − 1,*t*+1_), **z**_1,*t*+1_ = (*z*_1,*t*+1_, *z*_2,*t*+1_, …, *z*_0,*t*+1_), **z**_2,*t*+1_ = (*z*_2,*t*+1_, *z*_3,*t*+1_, …, *z*_0,*t*+1_, *z*_1,*t*+1_), and so on]; and (ii) the environmental state variables of all patches at the previous generation via **n**_**k**,*t*+1_. Due to the recursive nature of eq. (2), the environmental state variables in the focal generation, **n**_**0**,0_, depend on the whole history of traits **z**_H_ = (**z**_**0**,1_, **z**_**0**,2_, …) expressed in the population prior to the focal generation. As a result, the fitness of a focal individual may also depend on the traits expressed by all other previous individuals across space and time. To make this dependence explicit, we write the fitness of the focal individual as *w*(*z*_•_, **z**_**0**,0_, **n**_**0**,0_(**z**_H_)).

We make the additional assumption that in a monomorphic population where all individuals express the same resident phenotype *z*, the deterministic environmental dynamics described by the map *g* have a unique hyperbolically stable equilibrium point, identical in each patch, and satisfying

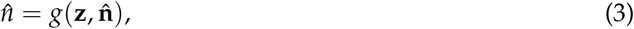

where **z** = (*z*, …, *z*) and 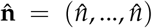 are vectors of dimension *D* whose entries are all equal to trait value *z* and environmental state variable 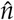, respectively. This is sometimes called the spatially homogeneous or flat solution in multi-patch ecological systems (p. 235 in [45]). The existence of such a solution entails that, in the absence of genetic variation, all patches converge to the same environmental equilibrium 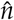, which may depend on the resident trait *z*. One useful property of a monomorphic population at such an equilibrium is that fitness must be equal to one, i.e. 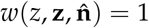 holds. This is because the total population size is constant, and consequently, each individual exactly replaces itself on average.

The fitness function (1) and the recursion (2) assume that fitness and the environmental dynamics can be written as functions of trait averages within patches. This said, this assumption does not limit us to only considering situations where effects within patches are additive. Indeed, because we are interested in convergence stability and thus in the first-order effects of selection (i.e. the first-order effects of trait variation), eqs. (1) and (2) are sufficient to model biological scenarios with arbitrary complicated non-additive traits effects among individuals within and/or between patches, for instance through complementarity or antagonism. Just a little care may be required when defining these expressions from an individual-based model (p. 95 in [5]).

### 2.3 Evolutionary dynamics

We assume that the quantitative trait evolves through rare mutations of small phenotypic effects, such that the evolutionary dynamics proceeds as a trait substitution sequence on the state space *Ƶ* ⊆ ℝ (i.e. the process of “long-term evolution” for finite populations described in [7]). We are interested in characterising convergence stable trait values, which are local attractors of the trait substitution sequence. To do so, we base ourselves on the first-order effects of selection on the fixation probability of a mutant that arises as a single copy in a population monomorphic for a resident trait value [11, 12, 7]. Technical details about our derivations can be found in S1 text and accompanying boxes. Our main findings are summarized below.

## 3 Results

### 3.1 Recipient-centered perspective: intra- and inter-temporal fitness effects

A trait value *z*\* ∈ *Ƶ* in the interior of *Ƶ* is convergence stable when

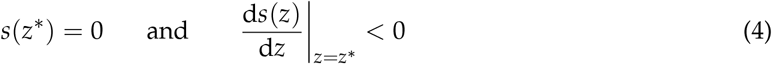

holds, where the function

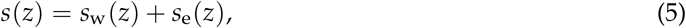

referred to as the selection gradient, can be written as the sum of two terms, given by

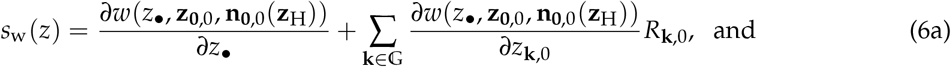

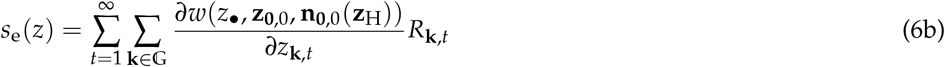

(see Appendix A in S1 Text for a derivation). Here, all quantities are evaluated in a monomorphic population with all individuals expressing the resident trait value *z*, and the environmental state variable in all patches are at the environmental equilibrium 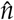 (eq. 3). A trait value *z*\* satisfying condition (4) constitutes a candidate end-point of evolution. More specifically, *z*\* is a mode of the stationary phenotypic distribution under the trait substitution sequence (see eq. A-5 and e.g. [7] for details). It is also possible for a mode of this distribution to lie on the boundary of a closed trait space, e.g. of *Ƶ* = [*z*_min_, *z*_max_] so that a mode sits at *z*_min_ and/or *z*_max_. For a mode to be at *z*_min_, *s*(*z*_min_) < 0 must hold, in which case *z*_min_ is convergence stable. Similarly, if *s*(*z*_max_) > 0 holds, then *z*_max_ is convergence stable and a mode of the stationary phenotypic distribution.

Both *s*_w_(*z*) and *s*_e_(*z*) depend on (i) marginal fitness effects, i.e. on derivatives of focal fitness (that we interpret below), and (ii) the relatedness *R*_**k**,*t*_ between the focal individual and another randomly sampled individual from patch **k**, *t*. Such relatedness coefficient is defined as

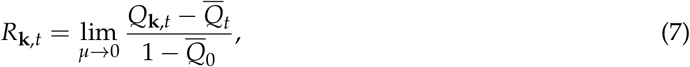

where *μ* is the mutation rate at the evolving locus; *Q*_**k**,*t*_ is the stationary probability that an allele sampled in the focal individual is identical by descent with a homologous allele sampled in another individual chosen at random from patch **k**, *t* under neutrality (i.e. in a population monomorphic for *z*); and 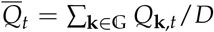 is the average probability of identity by descent between two homologous alleles sampled in two individuals living *t* generations apart. The probability of identity by descent *Q*_**k**,*t*_, and thus the relatedness coefficient *R*_**k**,*t*_, may depend on the resident phenotype *z*, but we leave this dependence implicit for readability.

Relatedness *R*_**k**,*t*_ quantifies the extent to which an individual that is sampled from patch **k**, *t* is more (when *R*_**k**,*t*_ > 0) or less (when *R*_**k**,*t*_ < 0) likely than a randomly sampled individual to carry an allele identical by descent to one carried by the focal individual at a homologous locus. To illustrate this notion, consider a Wright-Fisher process (where there is no adult survival and individuals are semelparous), which is the reference model for probabilities of identity by descent under isolation by distance (e.g. [46, 47, 5]). For this model, the relatedness coefficients *R*_**k**,*t*_ for *t* = 1, 2, 3, … can be shown to be given by

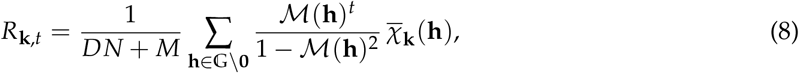

where 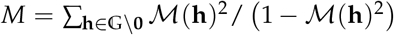 and ℳ(**h**) is the Fourier transform (or characteristic function) of the *m*_**k**_ probabilities (see Box 1 for definitions of Fourier transforms, character functions *χ*_**k**_(**h**), and their inverses 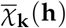 Appendix B in S1 Text for an example of the characteristic function of a dispersal distribution; and [48] for a derivation of eq. 8). The relatedness coefficient between two individuals in the same generation, *R*_**k**,0_, is given by eq. (8) with *t* = 2 (i.e. *R*_**k**,0_ = *R*_**k**,2_ holds for all **k**). In a panmictic or randomly mixing population (where *m*_**k**_ = 1/*D* for all **k**), relatedness between any two individuals is zero (i.e. *R*_**k**,*t*_ = 0 for all **k** and all *t*; as the Fourier transform reduces to *ℳ*(**h**) = 1 if **h** = **0** and 0 otherwise using property I.F in Box 1). However, as soon as dispersal is non-uniform (i.e. if *m*_**k**_ ≠ *m*_**j**_ for some **k** ≠ **j**), relatedness varies among individuals from different patches according to spatial and temporal distance. In particular, when dispersal is limited so that individuals have a tendency to remain in their natal patch, relatedness between the focal individual and individuals in the same patch from the same generation increases (*R*_**0**,0_ > 0). Because the average relatedness is zero (i.e. 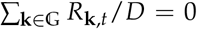 holds from eq. 7), the focal individual must also be negatively related to individuals residing in at least one other patch (i.e. *R*_**k**,0_ < 0 must hold for some **k** ≠ **0**). Which patches those are depends on patterns of dispersal. Under short-range dispersal, the focal individual tends to be positively related to individuals in patches nearby and negatively related to individuals further away (Fig 2C and Fig 3C). Under long-range dispersal, relatedness can be negative between individuals living in patches at intermediate distances (Fig 2D and Fig 3D).

**Figure 2:**
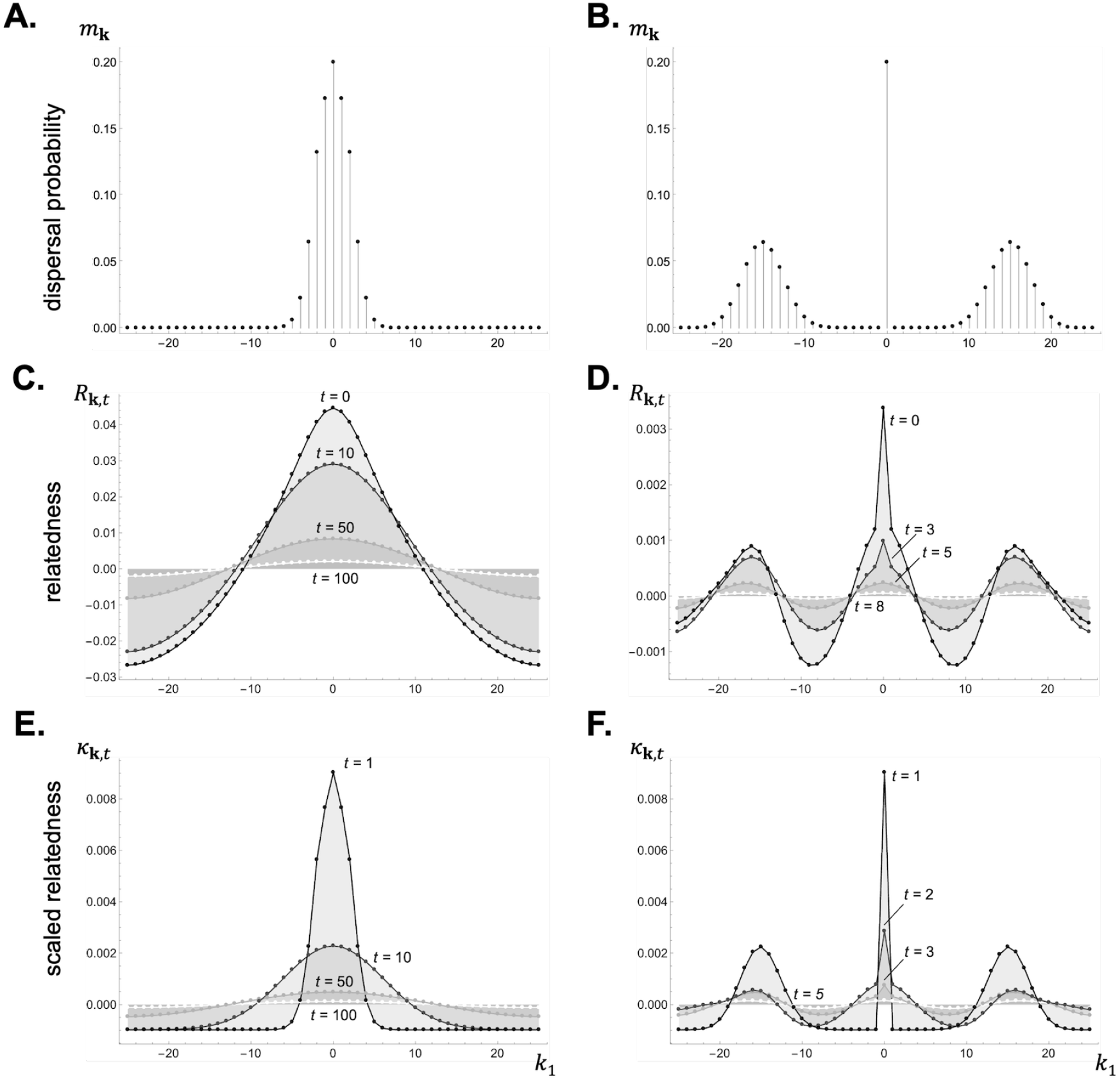
Dispersal distribution, relatedness, and scaled relatedness in a 1D lattice model under short and long-range dispersal. Panels **A-B**: Dispersal distribution *m*_**k**_ in a lattice-structured population in a one-dimensional habitat (with *D*_1_ = 51). An offspring leaves its natal patch with probability 1 − *m*_**0**_ = *m* = 0.8 and disperses to a patch at a Manhattan distance that follows a truncated binomial distribution (eq. A-7 in Appendix B.1 in S1 Text) with mean 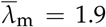 in panel A, leading to shortrange dispersal, and 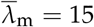 in panel B, leading to long-range dispersal. The distance dispersed along each habitat dimension is uniformly distributed across all dimensions and directions (Appendix B.1 in S1 Text for details). Panels **C-D:** Relatedness *R*_**k**,*t*_ for the dispersal distributions shown in panels A and B, respectively (using eq. 8 with patch size *N* = 20 and no adult survival s = 0). Panel C highlights how relatedness decays in time and space, becoming negative away from the focal deme when dispersal is short-range. In contrast, in panel D, where dispersal is long-range, relatedness is negative at intermediate and large distances, thus leading to a multimodal distribution of relatedness values. Panels **E-F:** Scaled relatedness *κ*_**k**,*t*_ for the dispersal distributions shown in panels A and B, respectively, under a Wright-Fisher model with fecundity effects (using eq. 21 with patch size *N* = 20). The trend of scaled relatedness is similar as that for relatedness. See S1 Data for how to generate these figures using Mathematica.

**Figure 3:**
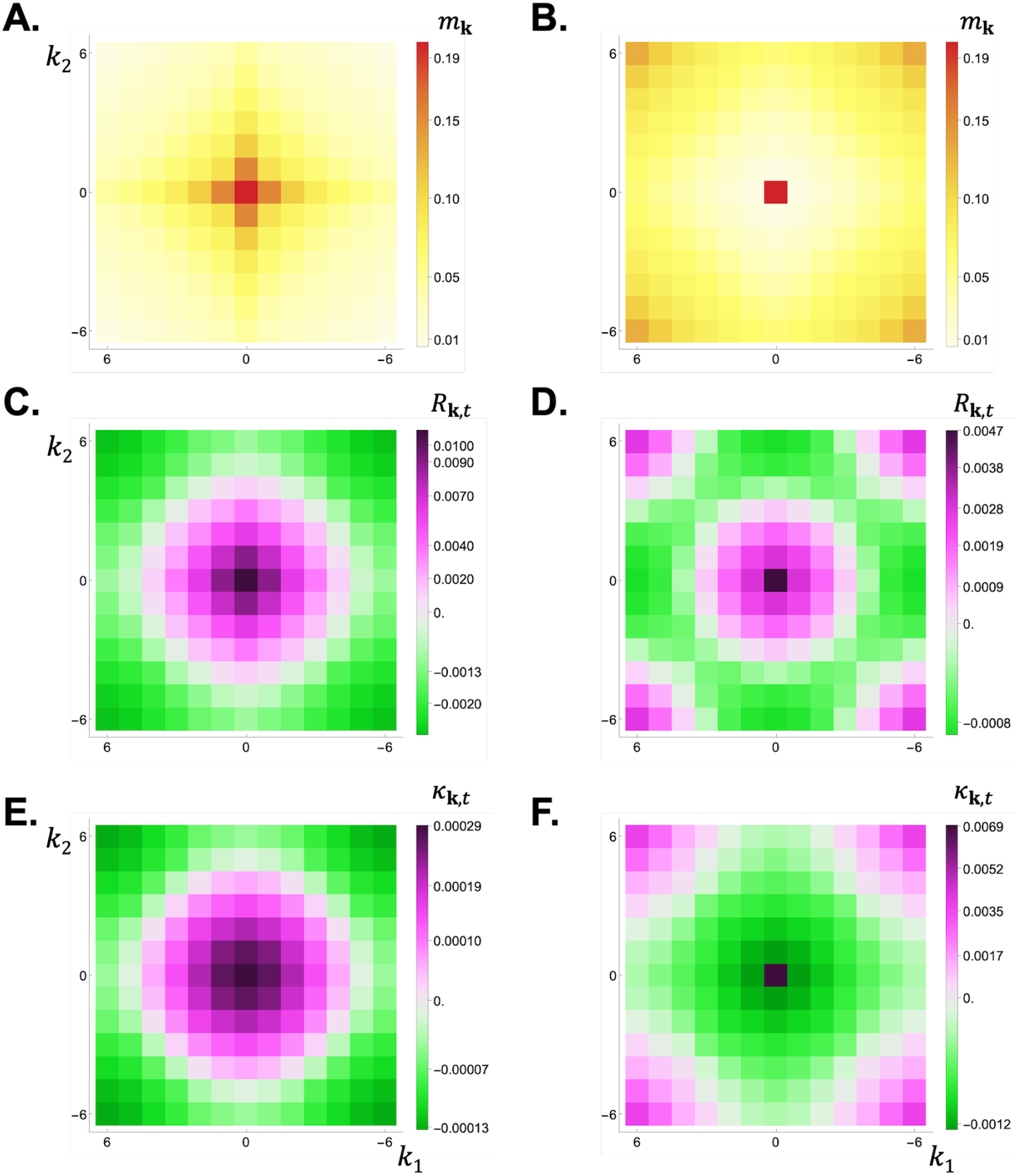
Dispersal distribution, relatedness, and scaled relatedness in a 2D lattice model under short and long-range dispersal. Panels **A-B:** Dispersal distribution *m*_**k**_ in a two-dimensional habitat (with *D*_1_ = *D*_2_ = 13). An offspring leaves its natal patch with probability 1 − *m*_**0**_ = *m* = 0.8 and <u>d</u>isperses to a patch at a Manhattan distance that follows a t<u>r</u>uncated binomial distribution with mean 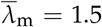 in panel A, leading to short-range dispersal, and 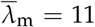 in panel B, leading to long-range dispersal (see Appendix B.2 in S1 Text for details). Panels **C-D:** Relatedness *R*_**k**,0_ from the dispersal distributions shown in panels A and B, respectively (using eq. 8 with patch size *N* = 20 and no adult survival s = 0). Panels **E-F:** Scaled relatedness, *κ*_**k**,10_ in panel E and *κ*_**k**,1_ in panel F, from the dispersal distributions shown in panels A and B, respectively, for a Wright-Fisher model (using eq. 21 with patch size *N* = 20). See S1 Data for how to generate these figures using Mathematica.

In eq. (6), the derivative *∂w*/*∂z*_•_ is the effect of a trait change in the focal individual on its own fitness. Likewise, *∂w*/*∂z*_**k**,*t*_ is the effect of a trait change in the whole set of individuals living in patch **k**, *t* on the fitness of the focal individual. Such effect is weighted by relatedness *R*_**k**,*t*_ in the expressions given by eq. (6). As such, *s*_w_(*z*) (eq. 6a) is the net effect of all intra-temporal effects on fitness, while *s*_e_(*z*) (eq. 6b) is the net effect of all inter-temporal effects (i.e. all effects within and between demographic periods, respectively). More broadly, eq. (6) consists in the sum of effects on the fitness of a focal individual stemming from all individuals that currently live (eq. 6a) or have lived (eq. 6b) in the population. Here, the focal individual is the recipient of phenotypic effects, present and past (recipient-centered perspective). How past phenotypic effects are mediated by environmental dynamics is left implicit in eq. (6), contained in eq. (6b) through **n**_**0**,0_(**z**_H_). In the next section, we expose such environmental effects by unpacking eq. (6b) and shifting from a recipient-centered to an actor-centered perspective.

### 3.2 Actor-centered perspective: environmentally mediated extended phenotypic effects

To understand natural selection on social traits, it is often helpful to see the focal individual as the actor, rather than the recipient, of phenotypic effects [49, 4, 5]. To shift to this perspective, we can leverage the space-time homogeneity of our model to see that *∂w*/*∂z*_**k**,*t*_ in eq. (6) is equivalent to the total effect of the focal individual on the fitness of the individuals in a patch at distance **k** at *t* steps in the future, and that relatedness *R*_**k**,*t*_ (eq. 8) quantifies the extent to which an individual sampled in a patch at distance **k** at *t* steps in the future is more (or less) likely than a randomly sampled individual to carry an allele identical by descent to one in the focal individual at a homologous locus [2]. These considerations readily lead to an actor-centered perspective for selection on intra-temporal effects, *s*_w_(*z*) (eq. 6a).

For selection on inter-temporal effects, *s*_e_(*z*), we further need to unpack the phenotypic effects through the environmental dynamics in eq. (6b). To do so, we now let the time index *t* ≥ 0 denote time forward so that **n**_**k**,*t*_ is the value of the environmental state variable in patch **k** at *t* time steps in the future of the focal generation (*t* = 0), and likewise let **z**_**k**,*t*_ denote the collection of population phenotypes at time *t* in the future. Environmental dynamics forward in time are characterised by rewriting eq. (2) as

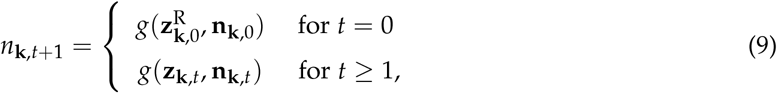

where 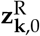 is equal to **z**_**k**,0_ except that the component *z*_**0**,0_ in this vector is replaced with 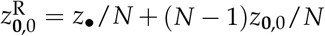, i.e. the average phenotype in the focal patch including the focal individual (e.g. for a one-dimensional lattice, 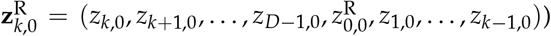. Eq. (9) brings upfront the potential complexity of characterising the environmental consequences of a trait change in the focal individual. This is because the trait of the focal individual, *z*_•_, influences the environmental state variable of potentially any patch **k** over one generation, *n*_**k**,1_, which can in turn have knock-on effects in the future on *n*_**k**,2_, *n*_**k**,3_, and so on throughout space in an interactive way. To characterise such effects, we write

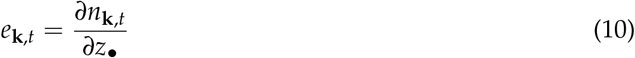

for the extended phenotypic effect of the focal individual on the environmental state variable in patch **k** at *t* generations in the future. Selection on inter-temporal effects *s*_e_(*z*) (eq. 6b) can then be written in terms of these extended phenotypic effects as

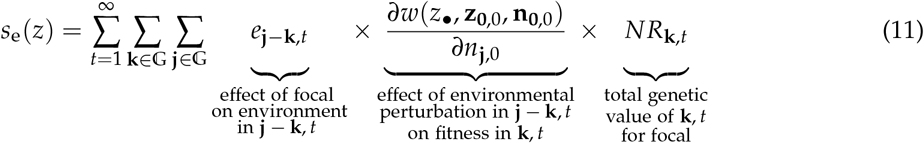

(see Appendix C.1 in S1 Text for a derivation). Eq. (11) indicates that *s*_e_(*z*) consists in the total effect of the focal individual on the fitness of individuals in each patch **k**, *t* in the future, via a change in the environmental state of possibly all patches **j** − **k**, *t*, where fitness is weighted by their relatedness *R*_**k**,*t*_ to the focal individual. From the point of view of the focal individual, relatedness *R*_**k**,*t*_ can then be thought of as the “genetic value” of an individual randomly sampled in patch **k**, *t* in units of fitness. More specifically, *R*_**k**,*t*_ can be interpreted as the number of units of its own fitness that the focal individual is willing to exchange with an individual from patch **k**, *t* against one unit of theirs without changing the mutant’s probability of fixation at *z*\*. Selection thus favours the focal individual sacrificing some of its own fitness to increase fitness in patch **k**, *t* when *R*_**k**,*t*_ > 0, and to decrease fitness when *R*_**k**,*t*_ < 0. How such sacrifice impacts the environment encountered by recipients is quantified by the extended phenotypic effect *e*_**j** − **k**,*t*_ in eq. (11).

The remaining challenge is how to compute *e*_**k**,*t*_, given the complex repercussions that a perturbation has in time and space (i.e. how to quantify a perturbation in the coupled dynamical system defined by eq. 9). We show in Appendix C.2 in S1 Text that this can be achieved through Fourier analysis using the following building blocks. First, we let *ψ*_**k**_ be the focal individual’s effect on the environmental state variable of patch **k** over one generation. Owing to our space-time homogeneity assumptions, this effect can be calculated as

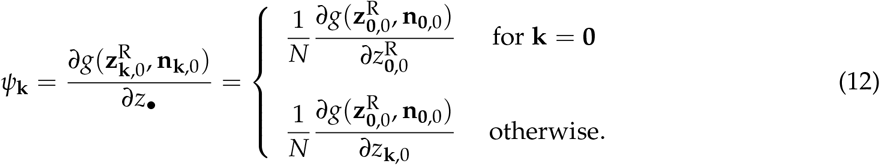

In practice, the rightmost expression is often more useful than the expression between equal signs in eq. (12) (as the the rightmost expression only requires characterising the environmental map 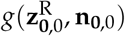 of the focal patch, e.g. eq. 26 below). Similarly, we let

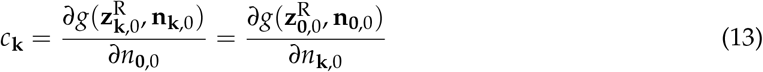

be the effect of the environmental state variable of one patch on the environmental state variable of another patch at distance **k** over one generation. With the above notation, and writing Ψ(**h**) and 𝒞(**h**) for the Fourier transforms of *ψ*_**k**_ and *c*_**k**_ respectively, the extended phenotypic effect can be efficiently computed as the inverse Fourier transform of 𝒞(**h**)^*t* − 1^Ψ(**h**) = ℰ_*t*_(**h**) (see Box 1 for the definition of inverse Fourier transforms), namely

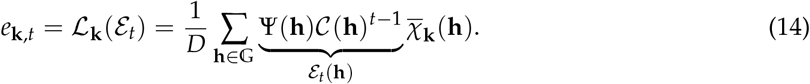

The form of ℰ_*t*_(**h**) indicates that *e*_**k**,*t*_ can be considered a perturbation in the dynamics of an environmental state variable that ripples into the future. This perturbation has its origin in a focal individual whose trait affects the environmental state variables of possibly multiple patches over one generation (captured by Ψ(**h**) in eq. 14). This one-generational change then propagates through space over *t* − 1 generations owing to the environmental dynamics, finally impacting the environment of individuals *t* generations downstream of the focal individual (captured by 𝒞(**h**)^*t* − 1^ in eq. 14). In Box 2, we generalize eqs. (11)-(14) to multi-dimensional environmental dynamics, that is, when multiple environmental state variables can be affected by the evolving trait and whose dynamics can interact with one another (e.g., metacommunity dynamics).

#### Box 2

**Multi-dimensional environment**

We generalize *s*_w_(*z*) and *s*_e_(*z*) to the case where there are *n*_e_ > 1 environmental state variables. We denote by 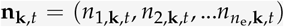 the vector of such variables in patch **k**, *t* (where *n*_*i*,**k**,*t*_ ∈ ℝ is the value of the *i*th environmental variable). The fitness of the focal individual is now given by

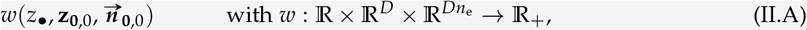

where 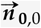 is the value at *t* = 0 of 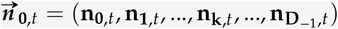, whose elements are solutions of

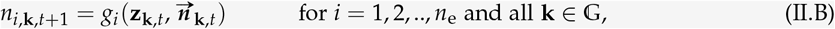

where *g*_*i*_ is the transition map for environmental variable *i*. We assume that in a monomorphic population *z*, there is a hyperbolically stable fixed point to environmental dynamics,

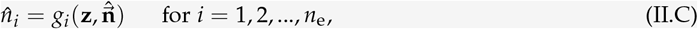

where 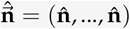 is a vector of dimension *D* whose entries are all given by 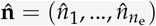. With fitness given by eq. (II.A), selection on intra-temporal effects *s*_w_(*z*) remains unchanged, given by eq. (6a) with 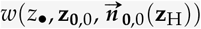 substituted for *w*(*z*_•_, **z**_**0**,0_, **n**_**0**,0_(**z**_H_))). For selection on inter-temporal effects *s*_e_(*z*), carrying out *mutadis mutandis* the same calculations as we have for the one-dimensional case, yields

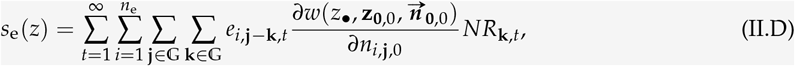

where *e*_*i*,**k**,*t*_ is the extended phenotypic effect on environmental variable *i* in patch **k**, *t*. This is computed as the inverse transform

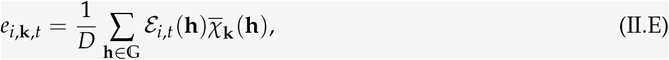

where ℰ_*i,t*_ (**h**) is the *i*-th element of the vector 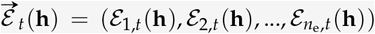, which is obtained from

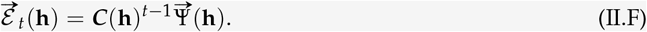

Here, the community matrix 𝒞(**h**) has its *ij*-th element given by 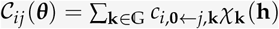, where

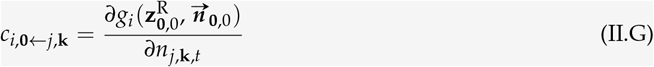

is the effect of environmental variable *j* in the focal patch on environmental variable *i* in patch **k**, *t*. The vector 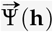 has *i*-th element given by Ψ_*i*_ (**h**), which is the Fourier transform of

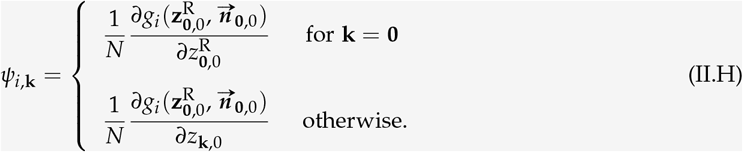

Under the infinite island model of dispersal, where *R*_**k**,*t*_ = 0, *e*_*i*,**k**,*t*_ = 0, and *c*_*i*,**0**←*j*,**k**_ = 0 for all 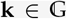 except **k** = **0**, eq. (II.D) reduces to eqs. 15-16 of [41].

Eq. (11) together with eqs. (12)-(14) constitutes a basis to quantify and understand selection on traits that have inter-temporal effects through the environment under isolation by distance. These equations formalise the intuition behind inclusive fitness arguments for environmentally mediated social interactions, saying that natural selection tends to favor traits whose environmental effects benefit the fitness of relatives (i.e. individuals more likely to carry genes that are identical by descent). Here, the fundamental currency is individual fitness, and its exchange rate among individuals is given by relatedness. However, in many cases it is not fitness that is directly impacted by traits or the environment, but rather some intermediate payoff, such as energy, the amount of prey caught, or the size of a breeding territory. In turn, this payoff influences survival and reproduction, which together determine fitness. We next explore such scenarios.

### 3.3 Payoff-mediated fitness: scaled relatedness or the genetic value of others in payoff units

#### 3.3.1 Payoff and fitness

Following much of evolutionary game theory (see e.g. [8] for a textbook treatment), we now consider cases where fitness depends on a payoff function summarizing social interactions (e.g. a case where the energy that an individual accrues depends on its foraging behaviour, the foraging behaviour of others, and how resources are distributed in the environment). We let this payoff function be *π* : ℝ× ℝ^*D*^ × ℝ^*D*^ → ℝ_+_, such that *π*(*z*_•_, **z**_**0**,0_, **n**_**0**,0_) is the payoff to the focal individual with phenotype *z*_•_ when the collection of average phenotypes among all other individuals is **z**_**0**,0_ and the collection of environmental state variables across all patches is **n**_**0**,0_. We assume that the fitness of this focal individual can, in turn, be written as a function 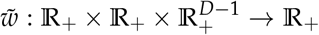 of the payoff to self, the average payoff to a patch neighbour, and the average payoff to an individual from each patch other than the focal, i.e. as

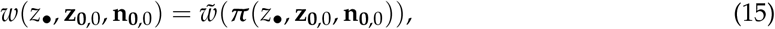

Where

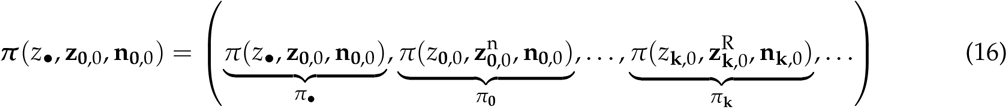

is a (*D* + 1)-dimensional vector collecting the payoff *π*_•_ to the focal individual, the average payoff *π*_**0**_ to a patch neighbour, and the average payoff *π*_**k**_ to an individual from each patch **k** ≠ **0**. As an argument to *π*_**0**_ in eq. (16), we used 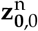 to denote a vector that is equal to **z**_**0**,0_ except for its first entry, which is given by 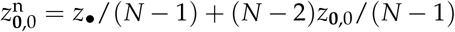, i.e. by the average trait among the neighbours of a neighbour of the focal individual. This captures the notion that the focal individual can influence the payoff of its neighbours.

Eq. (15) allows individual fitness to depend on the payoff of all the individuals of its generation in an arbitrary way. This said, in most applications the survival and fecundity of an individual depend only on its own payoff. In this case, fitness may be written as

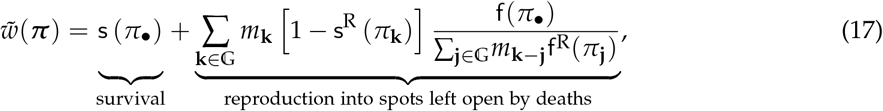

where s : ℝ_+_ → ℝ_+_ and f : ℝ _+_ → ℝ _+_ are survival and fecundity functions, respectively, and quantities with R as a superscript are defined so that s^R^(*π*_**0**_) = s(*π*_•_)/*N* + (*N* − 1)s(*π*_**0**_)/*N* is the average survival in the focal patch, otherwise s^R^(*π*_**k**_) = s(*π*_**k**_) for **k** ≠ **0**; and f^R^(*π*_**0**_) = f(*π*_•_)/*N* + (*N* − 1)f(*π*_**0**_)/*N* is the average fecundity in the focal patch, otherwise f^R^(*π*_**k**_) = f(*π*_**k**_) for **k** ≠ **0**. Suppose we set survival s to zero in eq. (17). In that case, we obtain the fitness function of the classical Wright-Fisher process (e.g. eq. 3 in [50], in the absence of environmental effects and for a circular stepping-stone model). More generally, if s is a positive constant and payoffs only influence fecundity f, eq. (17) implements a form of “death–birth” updating protocol (i.e. where individuals sampled at random to die are replaced by the offspring of selected individuals according to payoff, e.g. [36, 51]). Conversely, a “birth-death” updating is obtained by setting f to a positive constant and letting payoff influence survival s only. Eq. (17) will constitute a useful platform to explore more specific examples later, even though many of our results hold for the more general relationship between payoff and fitness given by eq. (15).

#### 3.3.2 Selection under payoff-mediated fitness

If fitness is of the form of eq. (15), the selection gradient can be written as

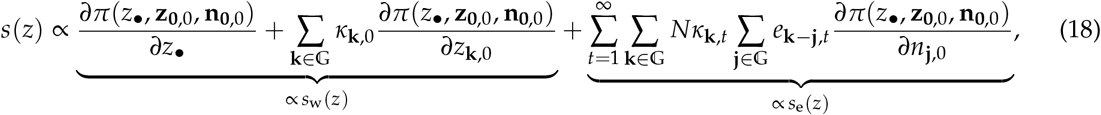

where

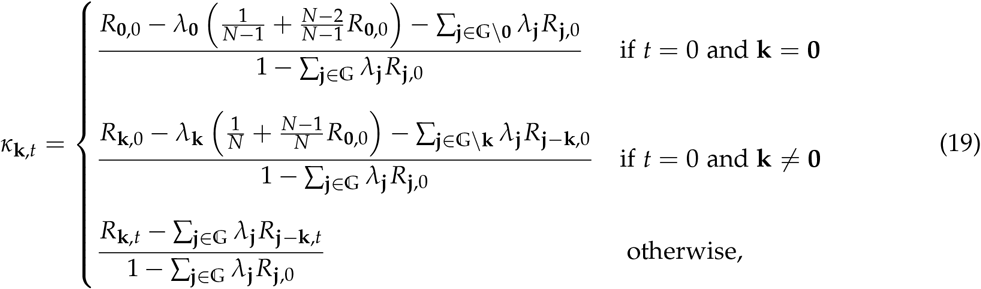

and

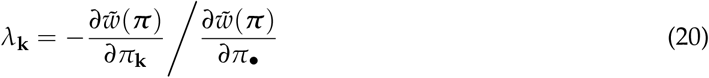

(see Appendix D in S1 Text for a derivation). To understand eq. (18), it is first useful to interpret *λ*_**k**_ (eq. 20) as a coefficient of fitness interdependence through payoffs. Specifically, *λ*_**k**_ measures the effect on the fitness of the focal individual of a change in the payoff of an individual at a distance **k**, relative to the effect of the payoff of the focal individual on its own fitness. When positive, *λ*_**k**_ can thus be interpreted as the strength of competition, as it indicates how much an increase in the payoffs of an individual at distance **k** reduces the fitness of the focal individual. With this in mind, the coefficient *κ*_**k**,*t*_ (eq. 19) can be thought of as a measure of relatedness scaled to competition (or scaled relatedness for short, e.g. [52, 53, 7]; with eq. 19 extending to isolation by distance the formalization of this concept found in [54]). To see this, suppose that the evolving trait helps current patch neighbours (i.e. such that *∂π*(*z*_•_, **z**_**0**,0_, **n**_**0**,0_)/*∂z*_**k**,0_ > 0) and consider the numerator of *κ*_**0**,0_ on the first line of eq. (19). This numerator consists in the relatedness *R*_**0**,0_ of a focal individual towards current patch neighbours, discounted by an inclusive fitness effect through increased competition, which is due to the increase in neighbours’ payoff caused by the focal individual helping them. The inclusive fitness effect consists of three relatedness weighted fitness effects: (i) − *λ*_**0**_/(*N* − 1), which captures the increase in competition experienced by the focal individual itself (where 1/(*N* − 1) is the frequency of the focal individual among the neighbours of an individual it helps); (ii) − *λ*_**0**_(*N* − 2)*R*_**0**,0_/(*N* − 1), which captures the increase in competition experienced by the relatives in the focal patch that are distinct from the individuals the focal individual is helping directly; and finally (iii) the total effect 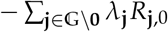, which captures the increase in competition experienced by relatives in other patches. The other two lines of eq. (19) are interpreted similarly: the relatedness towards neighbours from patch **k**, *t* discounted by an inclusive fitness effect on all recipients experiencing a change in fitness stemming from the focal changing the payoff of individuals in patch **k**, *t* (the denominator in eq. (19) standardizes these effects relative to the inclusive competitive effects induced by the focal individual changing its own payoff). More broadly, *κ*_**k**,*t*_ can be interpreted as the number of units of its own payoff that the focal individual is willing to exchange with a unit of payoff accruing to a randomly sampled individual from patch **k**, *t* without changing the mutant’s probability of fixation at a singular value *z*\*. The scaled-relatedness coefficient *κ*_**k**,*t*_ can thus be seen as the genetic value of other individuals in patch **k**, *t* relative to that of the focal individual in units of payoff. For (**k**, *t*) ≠ (**0**, 0), *κ*_**k**,*t*_ then measures the net social discount rate of temporally delayed and spatially extended fitness effects [43].

From the considerations above, eq. (18) can be read as an inclusive fitness effect at the payoff level. That is, selection depends on how the focal individual influences its own payoff and the payoff of all other individuals across patches, now and in the future, weighted by their scaled relatedness *κ*_**k**,*t*_. For recipients in the future (*t* ≥ 1), payoff effects are mediated by how the focal individual perturbs the environment in each patch (via *e*_**k** − **j**,*t*_ in eq. 18), and in turn by how such environmental perturbation influences payoffs (via *∂π*/(*∂n*_**j**,0_) in eq. 18).

In Appendix E in S1 Text, we use Fourier analysis to compute scaled relatedness *κ*_**k**,*t*_ for the fitness model given by eq. (17) and an arbitrary dispersal distribution *m*_**k**_. Our results are summarized in Box 3. For example, we obtain that under a Wright-Fisher process,

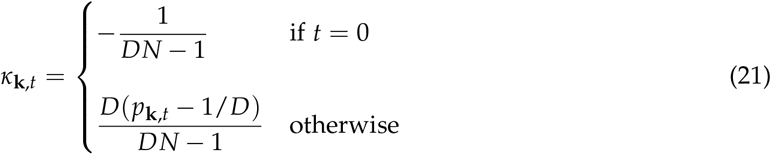

holds, where

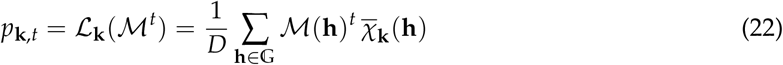

is the probability that, under neutrality, a gene descending from the focal individual is in patch **k** at *t* > 0 steps in the future (which depends on the characteristic function ℳ(**h**) of the dispersal distribution, Table 1). The collection of these probabilities, *p*_*t*_ = (*p*_**k**,*t*_)_**k**∈G_, can thus be seen as the distribution of a standard random walk on the set of patches with step distribution characterized by the dispersal probabilities *m*_**k**_. When such a random walk leads to a probability *p*_**k**,*t*_ that is greater than under a uniform distribution (i.e. when *p*_**k**,*t*_ > 1/*D* holds), eq. (21) indicates that scaled relatedness *κ*_**k**,*t*_ is positive, so that selection favours environmental transformations that increase payoffs in patch **k**, *t*. Conversely, selection favours environmental transformations that decrease payoffs in patches where *p*_**k**,*t*_ is smaller than under a uniform distribution (i.e. when *p*_**k**,*t*_ < 1/*D* holds). Which patches are those depends on the dispersal distribution (compare Fig 2E with Fig 2F for short and long-range dispersal in a 1D lattice, and Fig 3E with Fig 3F for short and long-range dispersal in a 2D lattice).

##### Box 3

**Scaled-relatedness coefficients. Scaled-relatedness coefficients**

With fitness given by eq. (17), we show in Appendix E in S1 Text that

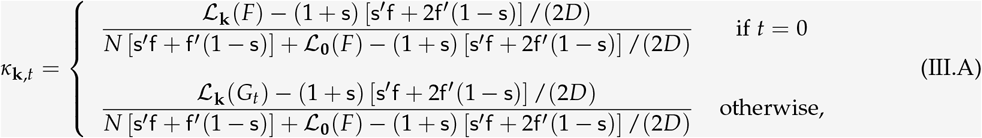

where as usual, all functions are evaluated at the resident trait value *z*, and ℒ_**k**_ (*X*) is the inverse Fourier transform of a function *X*(**h**) at 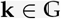 (eq. I.D of Box 1). The functions *F* and *G*_*t*_ are defined as

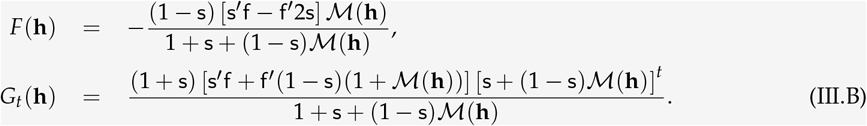

For fecundity effects under a Wright-Fisher process (s = s^′^ = 0), the above reduces to *F*(**h**) = 0 and *Gr*_*t*_ (**h**) = f^′^ ℳ(**h**)^*t*^, which yields eq. (21) of the main text when f^′^ = 1 (i.e. the payoff is directly fecundity). Using eq. (III.A), we also show in Appendix F in S1 Text that the summary statistic *K*, for selection on environmentally mediated social interactions under local interactions (eq. 24), is given by

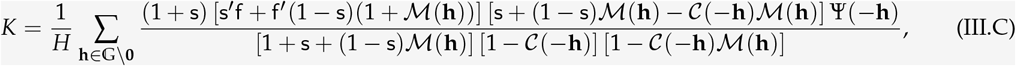

where

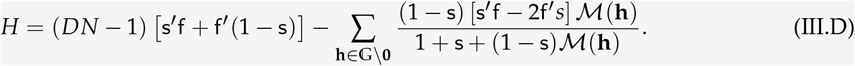

For fecundity effects under a Wright-Fisher process (s = s^′^ = 0) the summary statistic *K* simplifies to

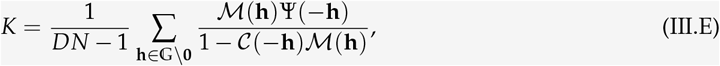

while for survival effects (f^′^ = 0), it simplifies to

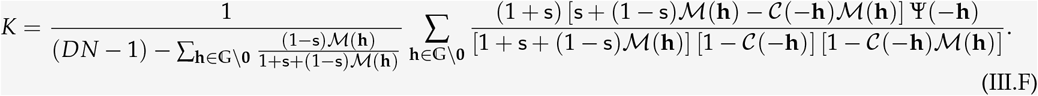

The selection gradient (18) recovers previous results of social evolution theory, which provides a robustness check of our calculations. To see these connections, assume that there are no ecologically meditated interactions, that is, that *∂π*(*z*_•_, **z**_**0**,0_, **n**_**0**,0_)/*∂n*_**0**,0_ = 0 holds, that − *C*(*z*) = *∂π*(*z*_•_, **z**_**0**,0_, **n**_**0**,0_)/*∂z*_•_ < 0 is a net payoff cost to self, and that *B*_**k**_(*z*) = *∂π*(*z*_•_, **z**_**0**,0_, **n**_**0**,0_)/*∂z*_**k**,0_ is a payoff benefit to individuals at distance **k**, which is typical of models under the heading of the evolution of “cooperation” or “altruism”. Further, suppose that individuals interact only with other individuals at a single distance **k** so that the selection gradient is proportional to − *C*(*z*) + *κ*_**k**,0_*B*_**k**_(*z*). Then, eq. (18) entails that selection favours such a helping behaviour if *κ*_**k**,0_ > *C*(*z*)/*B*_**k**_(*z*) holds, that is, if the scaled-relatedness coefficient is greater than the cost-to-benefit ratio. For a Wright-Fisher process, *κ*_**k**,0_ reduces to − 1/(*DN* − 1) ≤ 0 (eq. 21), which is always non-positive. Thus, helping cannot spread regardless of population structure because *κ*_**k**,0_ > *C*(*z*)/*B*_**k**_(*z*) cannot be satisfied as long as *B*_**k**_(*z*) > 0. This result was first derived for a lattice-structured population for *D* → ∞ and **k** = **0** in [55], and for finite *D* and any **k** under a circular one-dimensional habitat in [5] (chapter 8, and generalized to other abelian group structures in [44] and [43]). More generally, the scaled-relatedness coefficient given in Box 3 recovers established conditions for the spread of helping and harming behaviour in lattice-structured populations under different biological assumptions, such as for survival effects or for fecundity effects with overlapping generations (e.g. [56, 15, 36, 51]; see [43] for the explicit connections to this literature). Finally, in the presence of ecologically mediated interactions, so that *∂π*(*z*_•_, *z*_**0**,0_, *n*_**0**,0_)/*∂n*_**0**,0_ ≠ 0, eq. (18) recovers eq. (A.21) in [2] which holds for a Wright-Fisher process (to see this correspondence, set *s*_**k**,*t*_ = *N* (*∂π*(*z*_•_, *z*_**0**,0_, *n*_**0**,0_)/*∂n*_**0**,0_) *e*_**k**,*t*_ in eq. 18).

#### 3.3.3 Local interactions

In eq. (18), payoffs can depend on the traits expressed in and the environmental variables of all patches. In many instances, payoff effects can reasonably be assumed to be local, i.e. the payoff of an individual depends only on the traits and the environment of its patch. In this case, the payoff to the focal individual can be written as *π*(*z*_•_, **z**_**0**,0_, **n**_**0**,0_) = *π*(*z*_•_, *z*_**0**,0_, *n*_**0**,0_), and the selection gradient eq. (18) reduces to

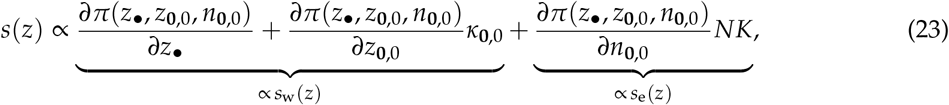

where

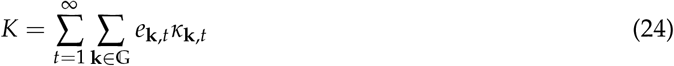

summarizes selection on environmentally mediated social interactions (see Appendix F in S1 Text for a derivation and eq. III.C in Box 3 for more details). When *K* = 0, selection is blind to the effects of the trait on the environment, even if the environment affects payoffs (i.e. even if *∂π*(*z*_•_, *z*_**0**,0_, *n*_**0**,0_)/(*∂n*_**0**,0_) ≠ 0). When *K* > 0, selection favours trait values that improve the environment (i.e. the payoff in the future increases). Conversely, when *K* < 0, selection favours trait values that deteriorate the environment (i.e. the payoff in the future decreases). Which of these outcomes unfolds depends on the interaction between extended phenotypic effects *e*_**k**,*t*_ and scaled relatedness coefficients *κ*_**k**,*t*_, with *K* > 0 when *e*_**k**,*t*_ and *κ*_**k**,*t*_ tend to be of the same sign (i.e. when 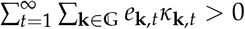 holds), and *K* > 0 when they tend to be of opposite sign (i.e. when 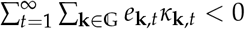 holds). In the next section, we explore through an example how this interaction depends on dispersal and how environmental state variables of different patches influence one another.

### 3.4 Inter-temporal helping and harming through a lasting commons

To gain more specific insights into how isolation by distance influences the way selection shapes environmentally mediated interactions, consider a scenario where the environmental variable is some lasting commons (e.g. a common-pool resource or a toxic compound) that can move in space, and whose production depends on an evolving trait that is individually costly to express. We assume that the commons is a “good” when the environmental variable takes positive values 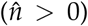 and a “bad” when it takes negative values 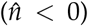. We also assume that the trait leads to the former when *z* > 0 and to the latter when *z* < 0, where the trait space is assumed to be *Ƶ* = ( − ∞, ∞).

The trait can thus be broadly thought of as environmentally mediated helping (increasing survival and reproduction to recipients) when *z* > 0, and as environmentally mediated harming (decreasing survival and reproduction to recipients) when *z* < 0.

We assume that fitness takes the form of eq. (17) with payoff given by

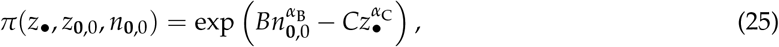

where *B* > 0 and *C* > 0 are parameters that respectively tune the effects of the environmental variable in the focal patch *n*_**0**,0_ and of the modifying trait *z*_•_ of the focal individual on the payoff of the focal individual. These effects also depend on the shape parameters *α*_B_ and *α*_C_, which we assume are positive integers, with *α*_B_ odd (e.g. *α*_B_ = 1) and *α*_C_ even (e.g. *α*_C_ = 2). Thus, the local commons increases (resp. decreases) payoffs when *n*_**0**,0_ > 0 (resp. *n*_**0**,0_ < 0) holds, but any trait expression, that is, any *z*_•_ away from 0, is individually costly and reduces individual payoff, and thus fitness. We also assume that costs increase more steeply than benefits, i.e. that *α*_C_ > *α*_B_ holds.

Meanwhile, how the trait modifies the commons is determined by the environmental map *g* (eq. 2), which we assume is given by

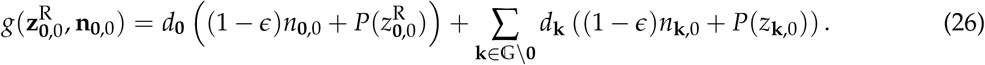

Eq. (26) states that the commons changes from one demographic time point to the next due to three processes. First, the commons is modified or “produced” in a patch according to a function *P* : ℝ→ ℝ of the average trait expressed in that patch, i.e. 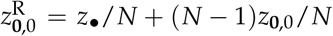 in the focal patch and *z*_**k**,0_ otherwise (with **k** ≠ **0**). We assume that *P*(0) = 0 holds, and that *P* is monotonically increasing, i.e. *P*′(*z*) > 0 for all *z* ∈ ℝ, where *P*′(*z*) denotes the derivative of *P* with respect to *z*. Second, each unit of commons “diffuses” or moves with probability *d*_**k**_ to a patch at a distance **k** from its source patch, where the probability distribution defined by *d*_**k**_ can be thought of as the environmental equivalent of the dispersal probability distribution *m*_**k**_. We let 𝒟(**h**) denote the Fourier transform of this distribution for future use (i.e. 𝒟(**h**) is to *d*_**k**_ what ℳ(**h**) is to *m*_**k**_, Table 1). Third, a unit of commons decays with rate 0 < *ϵ* ≤ 1 from one time step to the next. Substituting eq. (26) into eq. (3) indicates that in a monomorphic population for *z*, the dynamics of the commons stabilises to

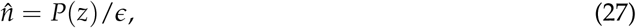

which is positive when *z* > 0, negative when *z* < 0, and whose absolute value increases as the rate of decay *ϵ* decreases, as expected. This equilibrium is unique because *g* is linear in *P*, and stable because *ϵ* > 0 holds.

With fitness of the form of eq. (17) and payoff eq. (25) only depending on local interactions (as in section 3.3.3), we can use eqs. (23) and (24) to characterise selection. With *κ*_**k**,*t*_ as given in Box 3, all that remains to be computed are the extended phenotypic effects, *e*_**k**,*t*_. To do so, we start by substituting eq. (26) into eqs. (12) and (13), to obtain *ψ*_**k**_ = *P*′(*z*)*d*_**k**_/*N*, and *c*_**k**_ = (1 − *ϵ*)*d*_**k**_, which in turn yield Ψ(**h**) = *P*′(*z*)𝒟 (**h**)/*N* and 𝒞(**h**) = (1 − *ϵ*) 𝒟(**h**). Substituting these expressions into eq. (14) then gives ℰ_*t*_(**h**) = (1 − *ϵ*)^*t* − 1^*P*′(*z*) 𝒟(**h**)^*t*^/*N*, which leads to

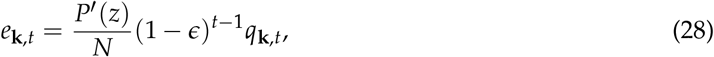

for the extended phenotypic effect *e*_**k**,*t*_ on patch **k**, *t*, where

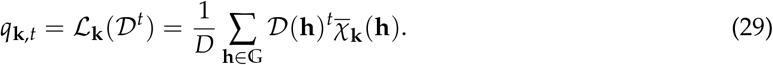

Equation eq. (28) can be understood as follows. By marginally changing its trait value, a focal individual produces *P*′(*z*)/*N* additional units of commons. Each such unit decays with time according to (1 − *ϵ*)^*t* − 1^, and ends up in patch **k**, *t* according to *q*_**k**,*t*_ (eq. 29), which can be interpreted as the probability that a non-decaying unit of the commons modified in the focal patch is located in patch **k** *t* generations in the future. In fact, the collection 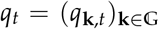 yields the distribution of a standard random walk on the set of patches with step distribution characterised by the *d*_**k**_’s. Extended phenotypic effects thus depend critically on the way the commons moves in space as captured by *d*_**k**_ (see Fig 4 for examples of *e*_**k**,*t*_ in a 1D model).

**Figure 4:**
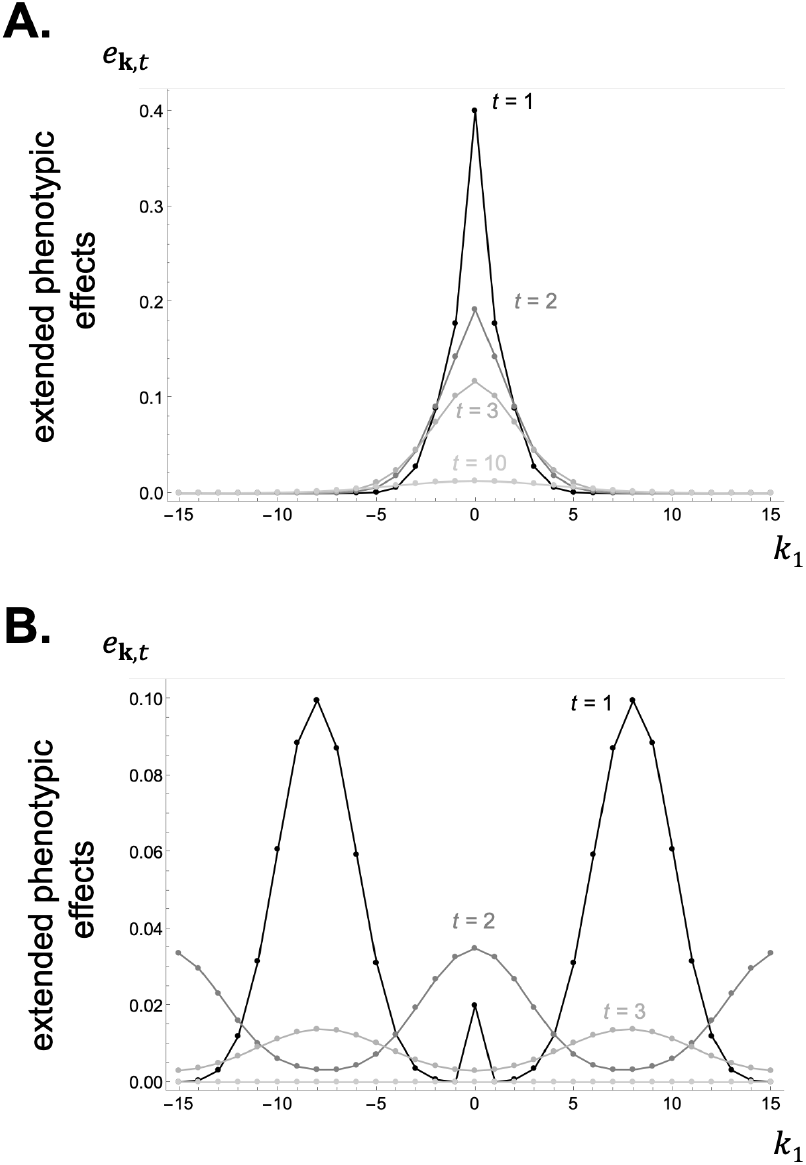
Extended phenotypic effects in a 1D lattice model under short and long-range movement of the commons. Panel **A**: When the commons moves locally, extended phenotypic effects *e*_**k**,*t*_ decay in time and space away from the focal deme (from eq. 28 with *D*_1_ = 31, movement probability *d* = 0.6 and expected movement distance 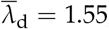, see Appendix G.5 in S1 Text for details on how movement is modelled; production function *P*(*z*) = *Nz*, i.e. e<u>a</u>ch unit of *z* contribute to one unit of resource; decay rate *ϵ* = 0.2; other parameters: *N* = 20, *m* = 0.3, 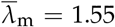). Panel **B**: In contrast, when the resource moves at greater distances, extended phenotypic effects *e*_**k**,*t*_ are greatest further away from the focal deme (from eq. 28 with movement parameters *d* = 0.98 and 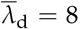 production function *P*(*z*) = *Nz*; decay rate *ϵ* = 0.5; other parameters: same as Fig 2A). See S1 Data for how to generate these figures using Mathematica.

In turn, how selection depends on extended phenotypic effects is found by substituting eq. (25) and eq. (28) into eq. (23). From this, we obtain

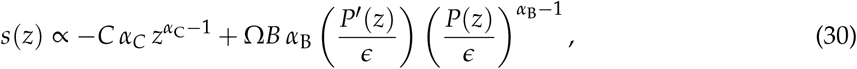

where

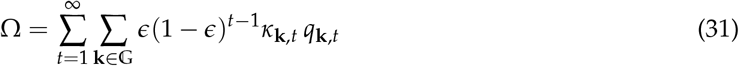

can be thought of as the total expected genetic value in units of payoff of all the individuals in the future that are affected by a unit of the commons in the focal patch. Since the term multiplying Ω in eq. (30) is positive (recall that *α*_B_ is odd and hence *α*_B_ − 1 is even), the selection gradient increases with Ω, and the greater Ω is, the greater the *z* favoured by selection. In fact, the selection gradient reduces to *s*(0) ∝ Ω at *z* = 0 (under our assumptions about parameters and the commons production function *P*). This shows that in a population where the trait is initially absent (so that individuals have no effect on the commons), selection favours environmental modifications leading to a common good (*z* > 0) when Ω > 0, or to a common bad (*z* < 0) when Ω < 0. Put differently, selection favours environmentally mediated inter-temporal helping when, in the eyes of the focal individual, the recipient of such help on average has positive genetic value in units of payoff, and conversely, inter-temporal harming when it has negative genetic value.

Further insights can be generated if we assume the simple functional form *P*(*z*) = *P*_0_*z*. In this case, we obtain

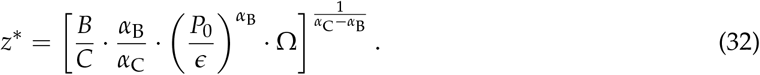

The singular value *z*\* (32) is convergence stable under our assumption that the cost of trait expression increases faster than the benefits (i.e. *α*_C_ > *α*_B_). Therefore, *z*\* is the mode of the stationary phenotypic distribution of the trait substitution sequence (we compute this distribution for the Wright-Fisher model in eq. A-136). From eq. (32) it is clear that the absolute value of *z*\* increases with the benefit-to-cost ratio *B*/*C*, with *α*_B_/*α*_C_, and with the environmental effect of the trait *P*_0_. However, whether *z*\* is positive or negative (and thus whether helping or harming evolves), ultimately depends on the sign of Ω, i.e. whether the expected genetic value Ω of a modification to the commons is positive or negative in payoff units.

The impact of species dispersal and commons movement on Ω can be understood most easily by assuming that payoff influences fecundity under a Wright-Fisher process (i.e. f^′^ > 0 and s^′^ = s = 0 in Box 3). In this case, Ω can be expressed as (see Appendix G.1 in S1 Text)

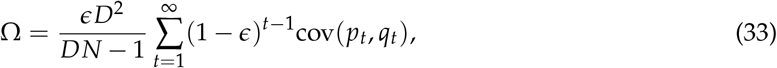

where cov(*p*_*t*_, *q*_*t*_) is the covariance between the distributions of the random walks of genes 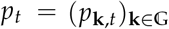 (eq. 22) and commons 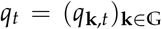 (eq. 29). This covariance is positive when there is a positive association between gene lineages and the commons these lineages modify. In other words, Ω is positive and helping is favoured when an environmental modification owing to the expression of a gene is most likely to be experienced by individuals living in the future and carrying identical-by-descent copies of that gene. Conversely, Ω tends to be negative and harming is favoured when this environmental modification is less likely to be experienced by future carriers.

While eq. (33) offers intuition on the biological conditions leading to positive or negative values of Ω, this quantity is more readily computed by noting that Ω = *ϵKN*/*P*^′^(*z*) holds, where *K* is defined in eq. (24), and by substituting eqs. (A-127) and (A-128) into eq. (III.E) in Box 3. Doing so, we obtain

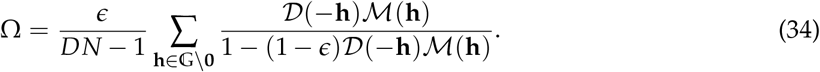

Fig 5A and Fig 5B give the sign of Ω under a binomial model for the distance of both the dispersal of the focal species and the movement of the commons (see Appendix G.5 in S1 Text for details). These figures show that such model of dispersal allows for both positive and negative values of Ω and thus for the evolution of both inter-temporal helping and harming. Here, helping corresponds to posthumous altruism and harming to posthumous spite since an individual can never obtain direct benefits from its own trait effects on the environment, and these effects are only felt by the next generation (as generations are non-overlapping under a Wright-Fisher process)

**Figure 5:**
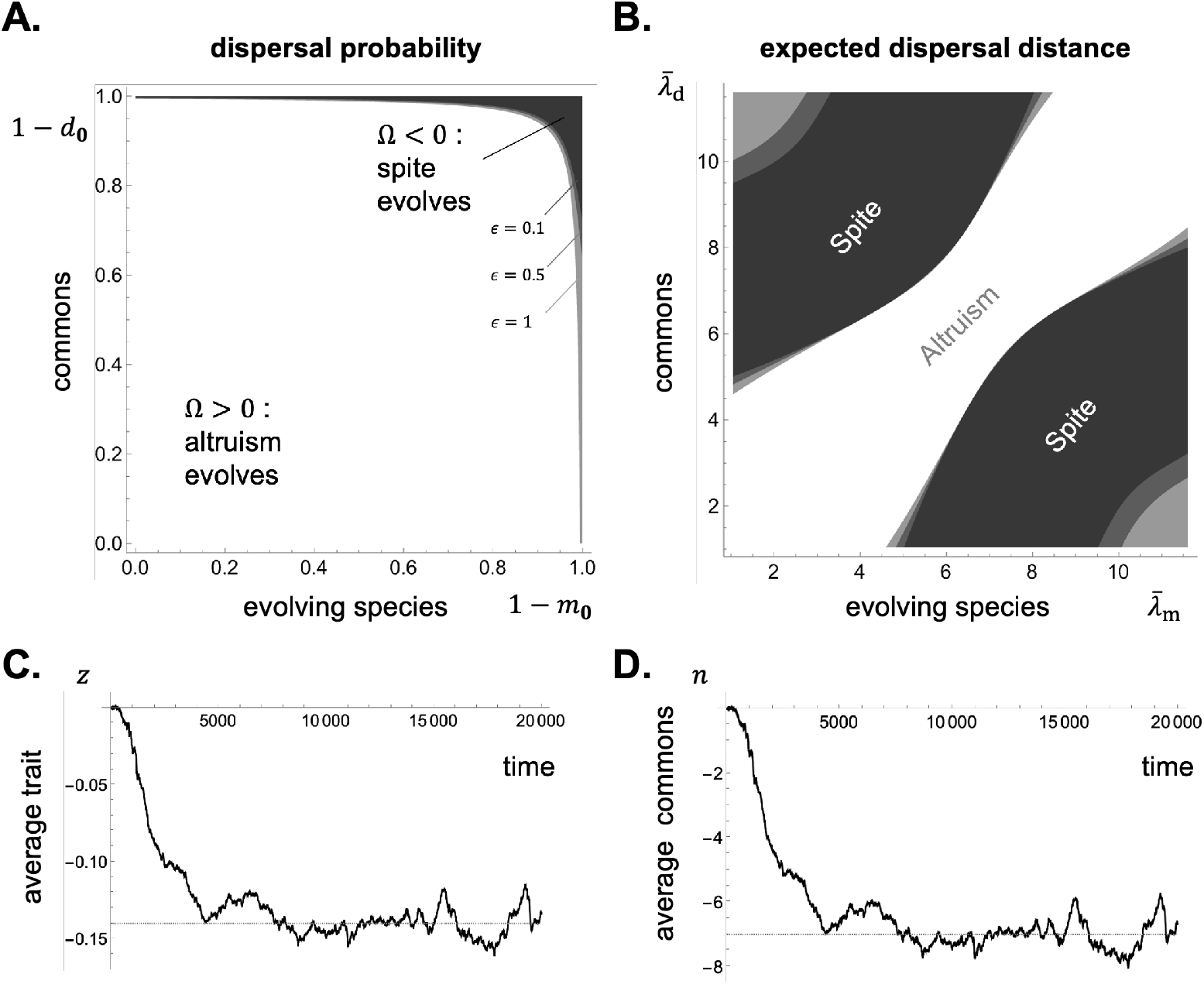
Selection favours altruism or spite depending on dispersal of the evolving species and the commons in a 2D lattice model. Panels **A**-**B**: Regions of dispersal parameters leading to the evolution of altruism, Ω > 0 (in white), or of spite, Ω < 0 (in gray) for an example in a 2D lattice model (with *D*_1_ = *D*_2_ = 13 and *N* = 50) under a Wright-Fisher life-cycle with fecundity effects (with Ω computed from eq. 34). Panel **A**: Combination of dispersal probability of the evolving species *m* = 1 − *m*_**0**_ (*x*-axis) and of the commons *d* = 1 − *d*_**0**_ (*y*-axis) for different levels of envir<u>on</u>mental decay *<u>ϵ</u>* in different shades of gray (*ϵ* = 0.1, 0.5, 1) with expected dispersal distance fixed (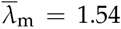 and 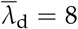). This shows that spite is favoured by high levels of dispersal <u>a</u>nd environmental decay. Panel **B**: Combination of expected dispersal distance of the evolving species 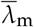(*x*-axis) and of the commons 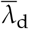(*y*-axis) for different levels of environmental decay *ϵ* in different shades of gray (see panel A for the legend) with dispersal probability fixed (*m* = 0.98 and *d* = 1). This shows that spite is favoured by dispersal asymmetry between the evolving species and the commons. Panels **C**-**D**: Evolution of spite in individual-based simula<u>ti</u>ons under a Wr<u>ig</u>ht-Fisher life-cycle with fecundity effects (with *D*_1_ = *D*_2_ = 13, *N* = 50, *m* = 0.3, 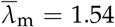, *d* = 1, 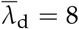, *B* = 2, *α*_B_ = 1, *C* = 1, *α*_C_ = 4, *P*(*z*) = *Nz*; for mutation, the trait mutates during reproduction with probability 10^−4^, in which case a normally distributed deviation with mean 0 and standard deviation 10^−2^ is added to the parental trait value). Panel C shows the average trait *z* in the population; panel D shows the average commons level or environmental variable *n* (with simulations in full lines – see S1 Code – and analytical prediction in dashed lines – from eq. 32 for *z* and 27 for *n*). See S1 Data for how to generate these figures using Mathematica.

More generally, numerical explorations of Ω (Fig 5A and Fig 5B) indicate that spite tends to be favoured by: (i) high levels of dispersal in the evolving species; (ii) high levels of movement of the commons; (iii) high environmental decay *ϵ*; and (iv) significant differences in the dispersal distance of the species and of the commons (e.g. when individuals disperse short distances but the commons move far away from their original patch). These conditions lead to a negative association between gene lineages and the commons these lineages modify. Conversely, altruism tends to be favoured when dispersal and movement are weak, environmental decay is low, and the distributions of the species’ dispersal and of the commons’ movement are similar (Fig 5A and Fig 5B, white region). In fact, under weak dispersal and movement (so that *m*_**0**_ = 1 − *m* and *d*_**0**_ = 1 − *d* with *m* and *d* close to zero), and regardless of the dispersal and movement distributions, we have

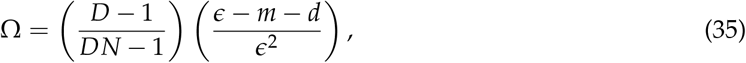

which is always positive when *m* and *d* are sufficiently small (see Appendix G.2 in S1 Text for a derivation of eq. 35). Under these assumptions, an individual’s lineage and the commons originating from its patch will be strongly and locally associated.

To see the effects of isolation by distance on social evolution, we compare in Fig 6A the singular strategy *z*\* for a 2D lattice where the evolving species and the commons follow a binomial model for distance travelled (detailed in Appendix G.5 in S1 Text), with the singular strategy *z*\* under the island model (where dispersal and commons movement are uniform). This comparison illustrates how isolation by distance allows for a wider range of evolved social behaviours as both altruism (*z*\* > 0) and spite (*z*\* < 0) can be favoured by selection depending on the distance dispersed. In contrast, only altruism is favoured by selection under the island model in Fig 6A (compare black full and dashed lines). Fig 6A also shows how selection may favour the evolution of more exaggerated social behaviours under isolation by distance (i.e. larger |*z*\*|). These differences are down to the fact that isolation by distance allows for a wider range in the covariance among genetic and commons movement (recall eq. 33). We also compared the singular strategy *z*\* found by substituting eq. (34) into eq. (32) with results from individual-based simulations. In these simulations, each offspring mutates with probability 10^−4^, in which case a normally distributed deviation with mean 0 and standard deviation 10^−2^ is added to the parental trait value. The only difference between these simulations and our analytical scenario is thus that multiple alleles can segregate due to mutation (rather than just two alleles under a trait substitution sequence, see [7] for finite populations). Nevertheless, we find an excellent fit between the convergence stable *z*\* and the mean population trait value in simulations (Fig 6B).

**Figure 6:**
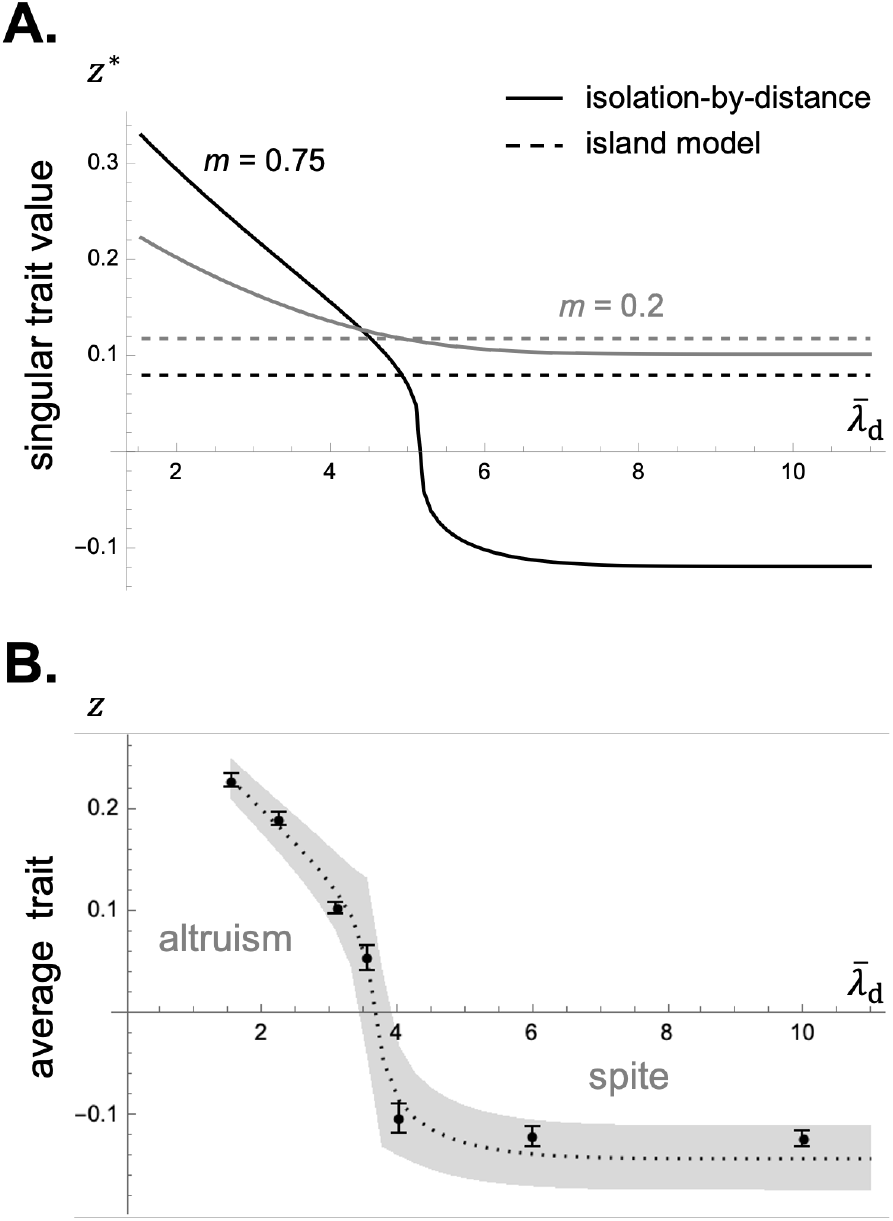
Isolation-by-distance allows for the evolution of a wider range of social behaviours than the island model under environmental feedback. Panel **A**: Singular trait value *z*\* from eq. 32 for *<u>m</u>* = 0.75 (in black) and *m* = 0.2 (in gray). Other parameters: *D*_1_ = *D*_2_ = 13, so that *D* = 169, *N* = 50, 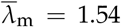, *d* = 0.99, *B* = 2, *α*_B_ = 1, *C* = 1, *α*_C_ = 4, *P*(*z*) = *Nz*. Dashed lines show the singular trait value with the same parameters but under the island model of dispersal for both the species and the commons (from eq. 32 with Ω given by eq. (A-126) derived in Appendix G.3). Panel **B**: Observed *vs*. predicted equilibrium trait value in individual-based sim<u>u</u>lations running for 20 000 generations under different expected dispersal distance of the commons *λ*_d_ leading to altruism (*z* > 0) and spite (*z* < 0). Other parameters: same as in Fig 5C and Fig 5D. The prediction is shown as a dashed line (from eq. 32) with grey region around for twice the standard deviation obtained from the stationary phenotypic distribution (from eq. A-136). Observed values of the trait average in the population are shown as black dots for the average from generation 5 000 to 20 000, with error bars for standard deviation over the same 15 000 generations. Simulations were initialised at the predicted convergence stable trait value. S1 Code for simulation code and S2 Data for simulation results.

The case where payoff influences survival rather than fecundity (s^′^ > 0 and f^′^ = 0 in eq. 17) is illustrated in Fig 7 (the expression of Ω for this case in terms of characteristic dispersal functions can be found in Appendix G.4 in S1 Text). This analysis reveals that harming tends to be favoured when baseline survival s is low, especially when environmental decay is also low (Fig 7C). This is because, otherwise, an individual may harm itself in the future. But apart from this, selection is not fundamentally different when payoff influences survival rather than fecundity in this model (i.e. under a birth-death *vs*. death-birth process).

**Figure 7:**
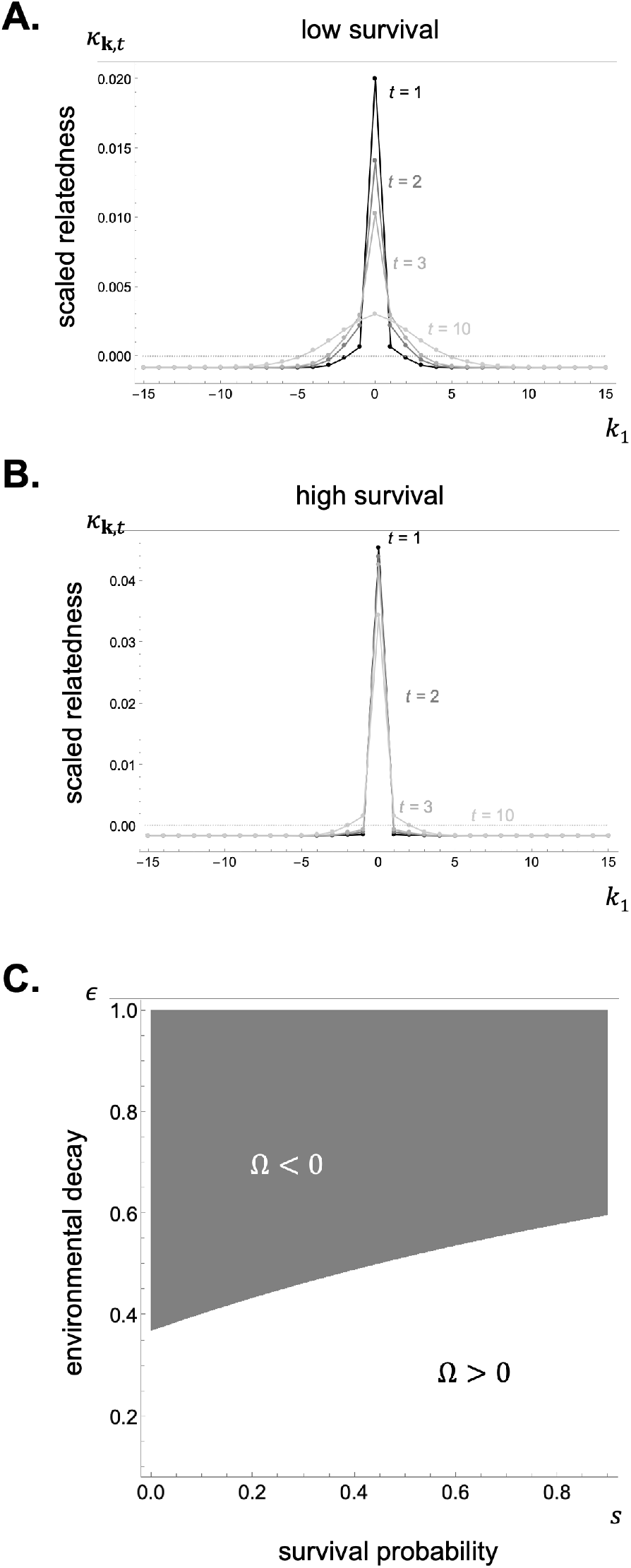
Scaled relatedness and selection under survival effects in a 1D lattice model. Panels **A-B:** Scaled relatedness *κ*_**k**,*t*_ in a 1D lattice model under survival effects (from Box 3 with s = 0 in panel A and s = 0.9 in panel B; other parameters: same as in Fig 4A). These panels show that genetic value decays away from the focal deme especially quickly when baseline survival is high (compare panels A and B). Otherwise, these profiles of scaled relatedness are similar to to those in Fig 4A, which suggests that selection acts similarly when the trait affects survival or fecundity. Panel **C**: Parameter region where selection favours the evolution of helping (Ω > 0) or harming (Ω < 0) under survival effects with adult survival probability s on the *x*-axis and environmental decay *ϵ* on the *y*-axis (Ω computed from eq. 31 using eq. A-129; other parameters: same as in Fig 4B, i.e. under long-range movement of the commons). See S1 Data for how to generate these figures using Mathematica.

## 4 Discussion

Our analyses characterise in two main ways the selection gradient on a trait that impacts the deterministic dynamics of environmental state variables that can be abiotic or biotic, which in turn feed back on individual survival and reproduction under isolation by distance.

First, we showed how selection on a trait due to its environmental effects can be understood in terms of how a focal actor influences the fitness of all future individuals via a modification to the environmental state variables that these individuals experience (eq. 11 and eq. II.D for the case of multiple environmental variables). The relevant trait-driven environmental modifications are formalized as extended phenotypic effects that quantify how a trait change in an actor individual in the present affects the environmental state variables in all patches at all future times (the *e*_**k**,*t*_ effects, eq. 14). While extended phenotypic effects are typically thought to benefit the actor or related contemporaries directly [57, 58], these effects in our model are all indirect, carrying over in space and time, thus influencing the fitness of future carriers of the actor’s trait when dispersal is limited. The associations between environmental and genetic variation that are necessary for selection to target trait-driven environmental modifications are given by the product between the extended environmental effects *e*_**k**,*t*_ and the relatedness coefficients *R*_**k**,*t*_ (see eq. 11), both of which can be efficiently computed using Fourier transforms (eq. 14 and II.E-II.F for a multivariate environment). These gene-environment associations indicate that selection favours traits or behaviours with environmental effects such that, when expressed by a focal individual, the environmental effects increase (resp. decrease) the fitness of future individuals that are more (resp. less) related to the focal than other individuals at that same future generation.

The second version of the selection gradient that we derived is based on the extra assumption that interactions between individuals are mediated through a payoff function (see eq. 17), as in most of traditional evolutionary game theory (e.g. [59, 60, 8]). Selection on a trait due to its environmental effects can still be viewed from an actor-centered perspective, but this time at the payoff rather than at the fitness level. Specifically, selection can be quantified in terms of how a focal individual influences the payoff of all future individuals via modifications to the environment these individuals experience, weighted by the relatedness between these individuals and the focal, now scaled to take competition into account (the term proportional to *s*_e_(*z*) in eq. 18, with scaled relatedness given in eqs. 19-20). The concept of scaled relatedness is useful because it summarizes in a single quantity (here one for each spatial and temporal distance) all the consequences of interactions among related individuals for indirect selection [61, 52, 53, 7]. That is, scaled relatedness balances, on the one hand, the positive effects of boosting the reproductive success of relatives in a particular spatial position, with, on the other hand, the negative effects of increasing competition for these relatives by affecting the reproduction and survival of others across the habitat. The increase of kin competition can be strong enough to offset the indirect benefits of social behaviour when social interactions occur among contemporaries (and generations do not overlap, e.g. [55, 5]). Because the strength of kin competition depends on the specifics of the life cycle (such as whether generations overlap or not, or whether payoff influences fecundity or survival), the evolution of direct social interactions is sensitive to such assumptions (see [52] for a review). This is notably the case under isolation by distance, where the evolution of altruism crucially depends on whether reproduction is modelled as a “death-birth” or a “birth-death” process (e.g. [62, 63, 64]). The “death-birth” process is akin to iteroparous reproduction with fecundity effects, and the resulting life cycle can sustain altruism; the “birth-death” process is akin to iteroparous reproduction with survival effects, and the life cycle inhibits altruism via increased kin competition (e.g. [56, 15, 17, 18]).

In contrast, we have found that in our model of environmentally mediated social interactions through a lasting commons, whether selection favours the evolution of helping or harming depends weakly on whether payoff influences survival or fecundity. There are two explanations for this. The first is that, because of environmental legacy, the effects on recipients are felt in the distant future, which decreases the competition among the focal’s own offspring [65, 41]. The second explanation is that, in our model, individuals and their environmental effects can move in space independently, further dissociating the positive and negative effects of interactions among relatives. This decoupling between benefits and costs means that natural selection can readily favor either posthumous altruism or posthumous spite in our model with non-overlapping generations. Which of these behaviours evolves depends on whether the combination of dispersal pattern and commons movement cause environmental effects to fall predominantly on individuals that are more or less related than average in the future.

Our findings on environmentally mediated posthumous spite merit further discussion as existing models suggest that the conditions for the evolution of spite are more restrictive than those for altruism. By spite, we refer here to a trait or behaviour whose expression decreases the individual fitness of both its actor and recipients. This is a strong form of spite, reducing the actor’s fitness (chapter 7 in [5]), which contrasts with the more commonly explored scenarios of weak spite (where the behaviour directly increases the actor’s fitness, e.g. [66, 67]). The evolution of strong spite typically relies on the existence of mechanisms by which individuals can evaluate their relatedness with social partners and thus behave according to some kin or type-recognition mechanism (e.g. [68, 69, 70, 71, 72]). By contrast, in our model spite is indiscriminate: an individual deteriorates the environment in the future without paying attention to recipients’ identities, even if this comes at a cost in the present. In the absence of generational overlap (e.g. Wright-Fisher model), such spite is entirely posthumous. With this in mind, it is noteworthy that spite can evolve even when local populations are not small (e.g. of local size 50, Fig 5). More broadly, our results illustrate how environmentally mediated social interactions under isolation by distance can evolve to be as relevant for fitness as direct social interactions.

The two main assumptions of our model are that fitness and environmental effects are homogeneous in space and time, and that environmental dynamics are deterministic. These assumptions are common to previous models interested in environmental or ecological changes in space that are evolutionarily driven, and particularly to those where individuals produce an environmental commons that moves according to a diffusion process (e.g. [73, 74, 75, 76, 77]). These models further assume a separation of time scales between demography and the commons such that in between the reproduction, death, or dispersal of any individual in the entire population, the dynamics of the commons reaches a stable distribution across the landscape. By contrast, here we have assumed that the mutation process, rather than the demographic process, is slow compared to environmental dynamics. Reproduction, death, or dispersal can occur on a similar time scale than environmental dynamics in our model, as is usually the case in ecological models (e.g. [78, 79]). As a result, even though environmental dynamics are described by a deterministic system (eq. 9, also as in most ecological models, [78, 79]), realised environmental dynamics fluctuate randomly on a similar time scale than unavoidable genetic fluctuations owing to finite patch size. The next challenge would be to consider a fully stochastic system for the environmental variables (i.e. to extend eq. 9 to the dynamics of a probability distribution). This would be especially useful to investigate the effects of demographic stochasticity in response to trait evolution [25], and to model, for instance, environmentally mediated evolutionary suicide or rescue. Our framework may nevertheless provide a suitable approximation to cases of demographic and environmental stochasticity (with eq. 9 giving the expectation in state variable at the next time step, conditional on the states of at the previous step). This approach has been shown to work well under the island model of dispersal provided that patches were not too small and dispersal not too limited [41]. It would be interesting to investigate how this holds up under isolation by distance. Finally, we considered the evolution of a single trait but our analyses apply directly to the case where multiple traits with environmental effects evolve jointly. To investigate such a case, one would first use our equations to obtain the selection gradient on each trait separately (applying eq. 5 to each trait under focus), and then jointly consider these gradients to perform a standard multidimensional analysis of directional selection (e.g. [80, 81]).

## Acknowledgements

The authors are grateful to Robbie I’Anson Price for help with Fig 1. CM thanks the Swiss National Science Foundation for funding, including salary (PCEFP3181243 to CM); JP acknowledges funding from the French National Research Agency (ANR) under the Investments for the Future (Investissements d’Avenir) program, grant ANR-17-EURE-0010, and from the Max Planck Institute for Evolutionary Anthropology.

## S1 Text

## Appendix A Convergence stability from fixation probability

Here, we prove eqs. (4)-(6) of the main text by considering the fixation probability of a single mutant with trait value *z* + *δ* into a population monomorphic for resident trait *z*. Let Π(*z* + *δ, z*) denote the fixation probability of this mutant, and by

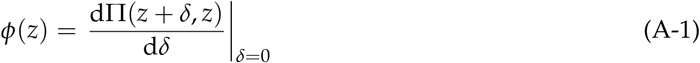

the derivative of the fixation probability with respect to the mutant effect. A trait value *z*\* that is convergence stable under a trait substitution sequence is thus characterized by

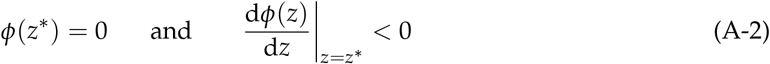

[1, 2, 3]. Under our modeling assumptions, the perturbation of the fixation probability is given by

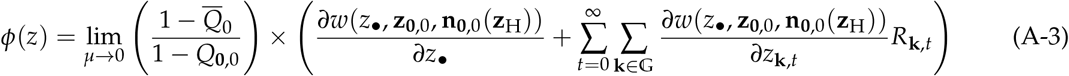

(eq. 1 of [4] together with eq. A11 of [5]), which can be expressed as

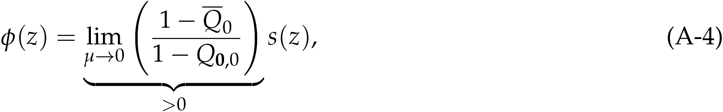

where *s*(*z*) is given by eqs. (5)–(6b). Because the limit in eq. (A-4) is always positive as long as *N* > 1 holds [2], the condition for convergence stability (A-2) is equivalently given by eq. (4).

The condition for convergence stability (A-2) also connects to the stationary probability density function *p*(*z*) that trait value *z* is observed in the population under a trait substitution sequence process in a finite population. This probability density function is given by

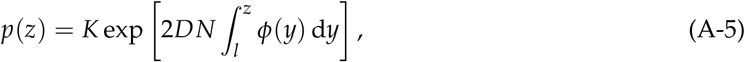

(eq. 7. of [6], eq. 62 of [3]) where *l* is the lower boundary of the state space and *p*(*z*) has a local maximum at *z*\* if conditions (A-2) are satisfied (see e.g. [6, 3] for details). The density function (A-5) is useful to evaluate the expected phenotypic variance in the population and can thus be compared to results from individual-based simulations (see eq. A-136 and Fig 6B for a concrete example).

To compute the probability density function *p*(*z*), however, requires to fully quantify the derivative of the fixation probability *ϕ*(*z*), which in turn depends on 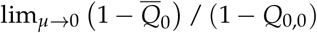, which is process specific. For instance, for the Wright-Fisher process,

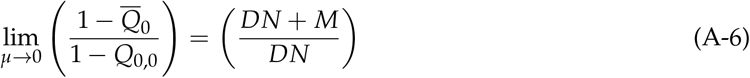

holds (eq. A17 in [4]), where 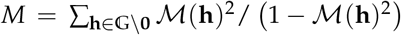 is defined as under eq. (8) in the main text and may remain complicated to evaluate. Yet evaluating eq. (A-6) may actually not be needed to compute *ϕ*(*z*). For instance when the selection gradient takes the form of eq. (A-55) (eq. (19) of the main text), eq. (A-6) cancels from *ϕ*(*z*) when fitness takes the form eq. (17) owing to eq. (A-105) (and this property may hold more generally). Further, using coalescent arguments, eq. (A-6) may be computed indirectly. For instance, owing to eq. (3.68) and eq. (3.70) of [2], one actually has 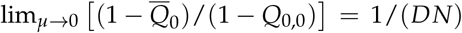 for the Wright-Fisher process under the infinite allele model.

## Appendix B A distribution for short and long range dispersal

Here, we specify a dispersal distribution based on the binomial distribution, which allows us to consider both short and long dispersal, and that we used to generate the various numerical examples of our analysis.

### Appendix B.1 One-dimensional habitat

Let us first consider a one-dimensional habitat consisting of a circular lattice, so that the set of patches is 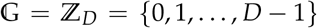, i.e. the set of integers modulo *D*. We assume that *D* is odd, so that we can write ℤ_*D*_ = {0, 1, …, (*D* − 1)/2, − (*D* − 1)/2, − (*D* − 1)/2 + 1, …, − 1}. We further assume that an individual disperses with probability *m*, and that it stays in its natal patch with probability 1 − *m*. If an individual disperses, it does so with equal probability either “clockwise” or “counterclockwise” a number *j* ∈ {1, 2, …, (*D* − 1)/2} of steps, which we assume follows a zero-truncated binomial distribution with probability mass function

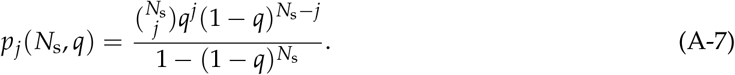

Here, *N*_s_ = (*D* − 1)/2 is the number of trials, and *q* = 2*λ*_m_/(*D* − 1) is the probability of success, where *λ*_m_ = *N*_s_*q* is the mean of the non-truncated distribution. The mean number of steps an individual disperses conditional on dispersal is given by

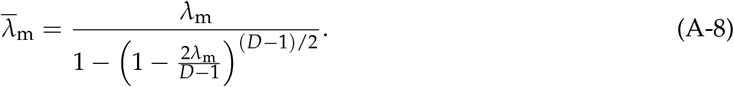

From these assumptions, the dispersal distribution is given by

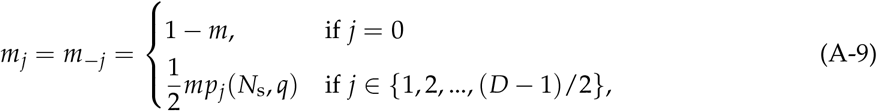

and its associated characteristic function (or Fourier transform I.B) can be written as

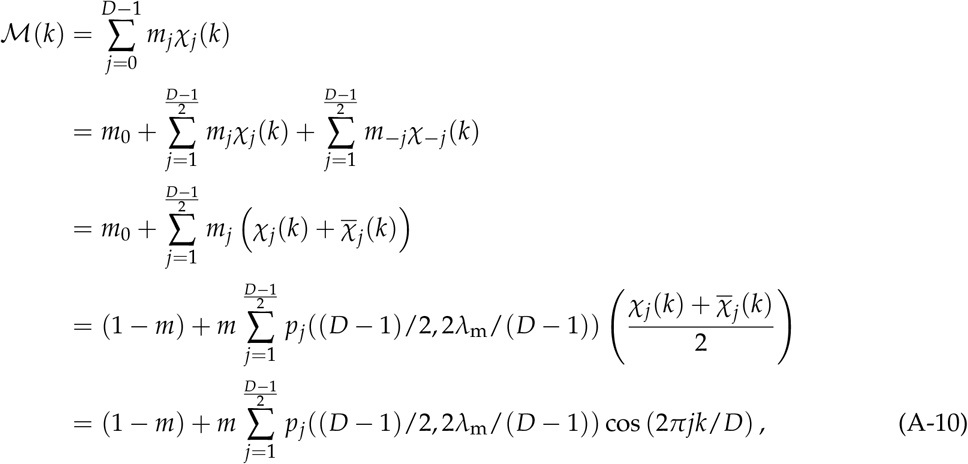

where the third line uses the fact that the migration kernel is symmetric (*m*_*j*_ = *m*_−*j*_ holds for *j* ∈ {1, 2, …, (*D* − 1)/2}) and the identity 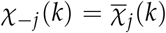, and the last line uses the trigonometric identity cos(*x*) = (exp(*ιx*) + exp( − *ιx*)) /2. Eq. (A-10) shows that the characteristic function of the dispersal distribution is determined by the parameters *D, m*, and *λ*_m_.

### Appendix B.2 Two-dimensional habitat

For the two-dimensional case, we consider a torus with the same number of patches in each dimension so that 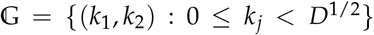 for *k*_1_ and *k*_2_ modulo *D*^1/2^. The dispersal distribution of the focal species *m*_**k**_ for 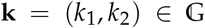, is constructed similarly as above. First, an individual disperses with probability *m* and with probability 1 − *m* stays in its natal patch. Second, conditional on dispersal, we sample the number of steps *j* ∈ {1, 2, …, *D*^1/2^ − 1} an individual disperses on the lattice (maximum *D*^1/2^ − 1) from a zero-truncated binomial distribution *p*_*j*_(*N*_s_, *q*) (eq. A-7) with parameters *N*_s_ = *D*^1/2^ − 1 and *q* = *λ*_m_/(*D*^1/2^ − 1). Accordingly, the mean number of steps an individual disperses conditional on dispersal is

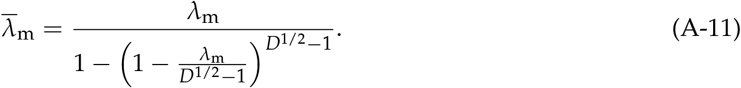

Third, we determine how this total number of steps *j* is divided between *j*_1_ steps in dimension 1 and *j*_2_ steps in dimension 2 (so that *j* = *j*_1_ + *j*_2_), assuming that dispersal in either dimension has the same distribution. We do so by sampling *j*_1_ from a discrete uniform distribution unif(*j*_min_, *j*_max_), where

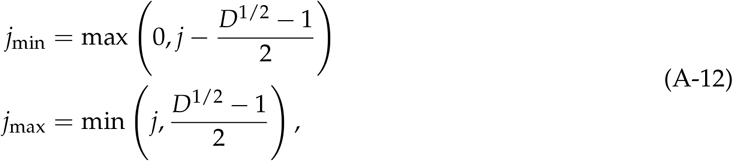

and by setting *j*_2_ = *j* − *j*_1_. Finally, given the number of steps in each dimension *j*_1_ and *j*_2_, these are then equally likely to occur in either direction away from the focal patch.

## Appendix C Extended phenotypic effects

### Appendix C.1 Actor-centered representation of inter-temporal effects

Here, we derive eq. (11) of the main text. To this end, we first apply the chain rule to the fitness expression (1) whereby we have for *t* ≥ 1 that

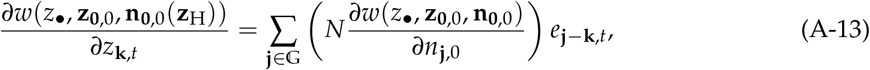

where we have defined

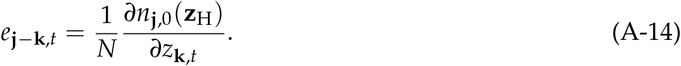

Thanks to spatial homogeneity this is in turn equivalent to

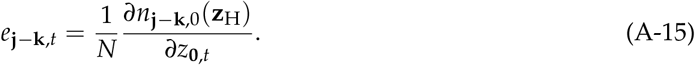

The quantity *e*_**j** − **k**,*t*_ is the extended phenotypic effect of a single individual residing in the focal patch at *t* time steps in the past on the value of the environmental variable in patch **j** − **k** in the present, where *∂n*_**j** − **k**,0_(**z**_H_)/*∂z*_**0**,*t*_ is the effect of the whole set of individuals in the focal patch at *t* time steps in the past on the value that the environmental variable takes in patch **j** − **k** in the present. But since the map *g* (eq. 2) does not depend on time (i.e. environmental dynamics are homogeneous in time), *e*_**j** − **k**,*t*_ is also the effect of a focal individual residing in the focal patch on the value that the environmental variable takes in patch **j** − **k** at *t* time steps in the future. We can thus write

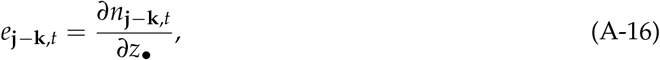

where *n*_**k**,*t*_ now stands for the value of the environmental variable in patch **k** at *t* steps in the future. Substituting eq. (A-16) into eq. (A-13), and this into eq. (6b), obtains eq. (11), as required.

### Appendix C.2 Extended phenotypic effects

Here, we derive the expression for the extended phenotypic effect given by eq. (14) of the main text. To do this, we first take the derivative on both sides of eq. (9) with respect to *z*_•_, which yields

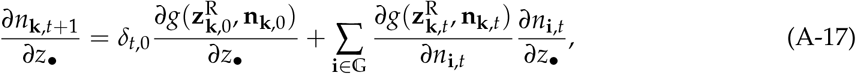

where *δ*_*t*,0_ is a Kronecker delta, and where we used 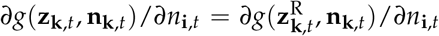, since all derivatives are evaluated at *z* and 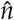. This also entails that the derivatives of the transition map *g* are independent of time, which allows us to write

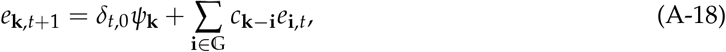

with

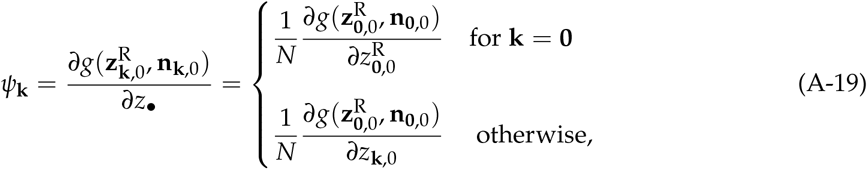

and

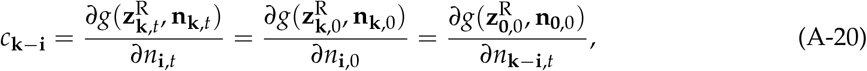

where the second equality in equation (A-19) follows from spatial homogeneity and the chain rule of derivatives, the second equality in equation (A-20) follows from temporal homogeneity, and the last equality in equation (A-20) follows from spatial homogeneity. These expressions are useful in concrete applications since only 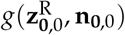 needs to be specified to evaluate *ψ*_**k**_ and *c*_**k**_ (see section 3.4).

We can solve eq. (A-18), using the Fourier transforms (see Box 1) 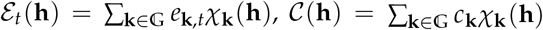 (Table 1) and 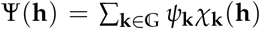 (Table 1). Using these expressions, from (A-18), and noting that *χ*_**k**_(**h**) = *χ*_**k** − **i**_(**h**)*χ*_**i**_(**h**) holds, we have

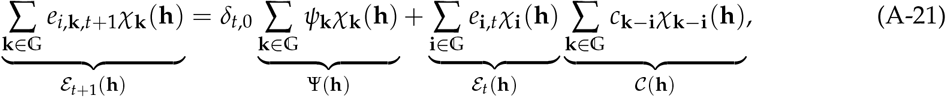

where the expression for 𝒞(**h**) holds by changing the dummy index of the sum. Thus, we obtain the recursion

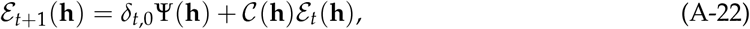

whose solution given the initial condition ℰ_0_(**h**) = 0 (as there are no extended phenotypic effects in the focal generation) is ℰ_*t*_(**h**) = 𝒞(**h**)^*t* − 1^Ψ(**h**), as required in eq. (14).

## Appendix D Selection gradient in terms of scaled relatedness

Here, we derive eq. (18) which, recall, is premised on the fitness of the focal individual taking the form

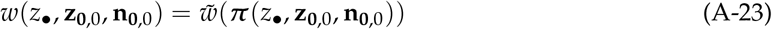

with payoff vector

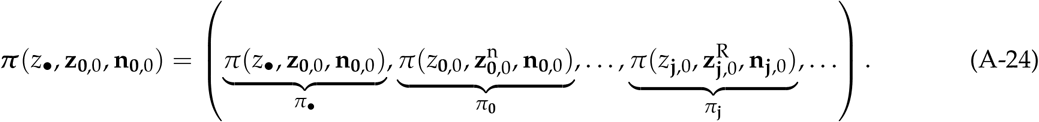

Here, 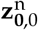 is equivalent to **z**_**0**,0_ except for the first entry which is given by

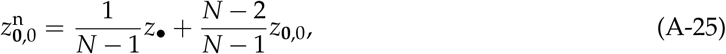

(instead *z*_**0**,0_ of in **z**_**0**,0_), and 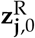 is equal to **z**_**j**,0_ except that the entry with component *z*_**0**,0_ in this vector is replaced with

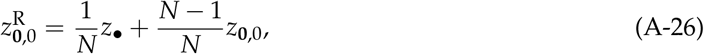

that is, with the average phenotype in the patch **0**, 0 including the focal individual.

To simplify the operation of taking derivatives of fitness with respect to phenotypes later, we first express the derivatives of the payoff *π*_**j**_ appearing in eq. (A-24) with respect to its various arguments in terms of the derivatives of the payoff to the focal individual. Applying the chain rule of derivatives and evaluating the derivatives at the resident phenotype, we readily obtain the following,

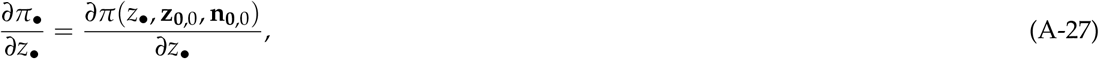

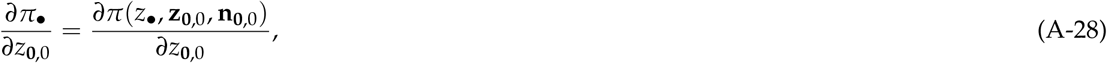

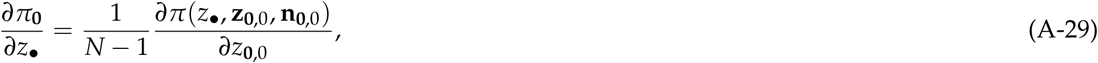

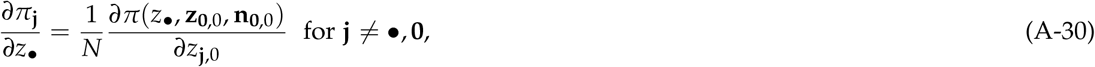

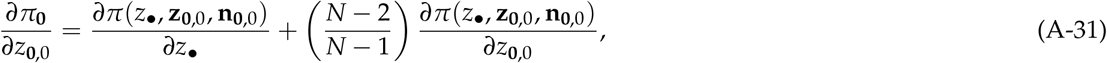

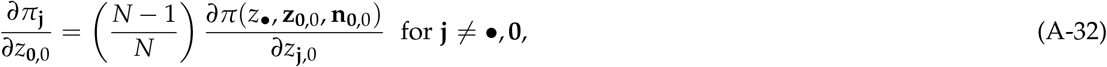

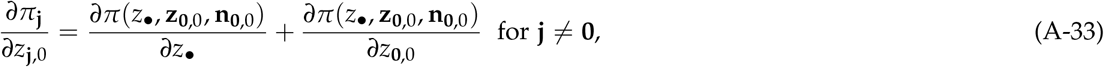

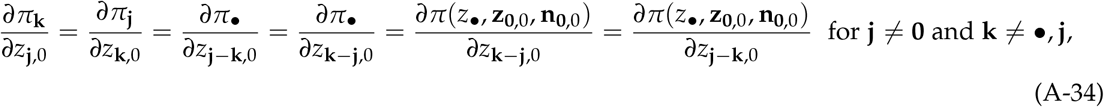

where the equalities in the last expression all follow from our assumption of spatial homogeneity.

Similarly, for derivatives of payoffs with respect to environmental state variables, we have

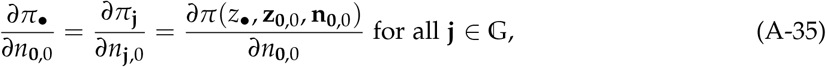

where the first and second equalities are consequences of spatial homogeneity, and

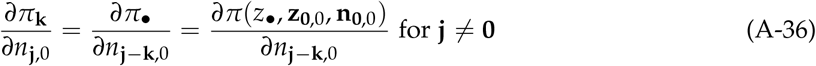

where the first equality is again a consequence of spatial homogeneity.

We can then write the derivatives of fitness that appear in the selection gradient (eqs. 6a–6b) in terms of the derivatives of the payoff to the focal individual (eqs. A-27–A-36) by applying the chain rule of derivatives to the right-hand side of eq. (A-23) and simplifying, as follows. First, the fitness derivative with respect to the focal individual’s phenotype can be written as

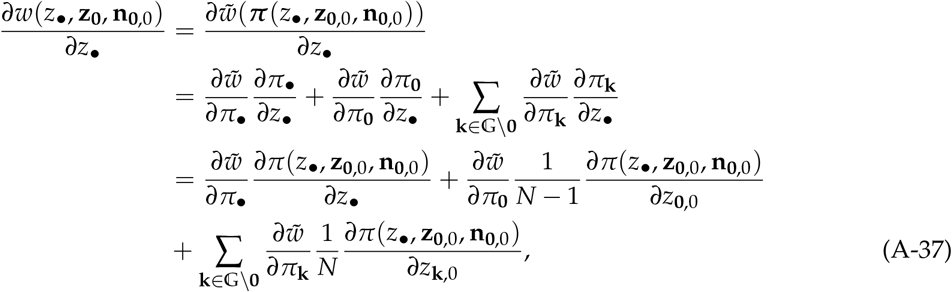

where the first equality follows from taking the derivative to both sides of eq. (A-23); the second equality follows from applying the chain rule; and the third equality follows from substituting eqs. (A-27)–(A-30).

Second, the fitness derivative with respect to the average phenotype of patch neighbours is

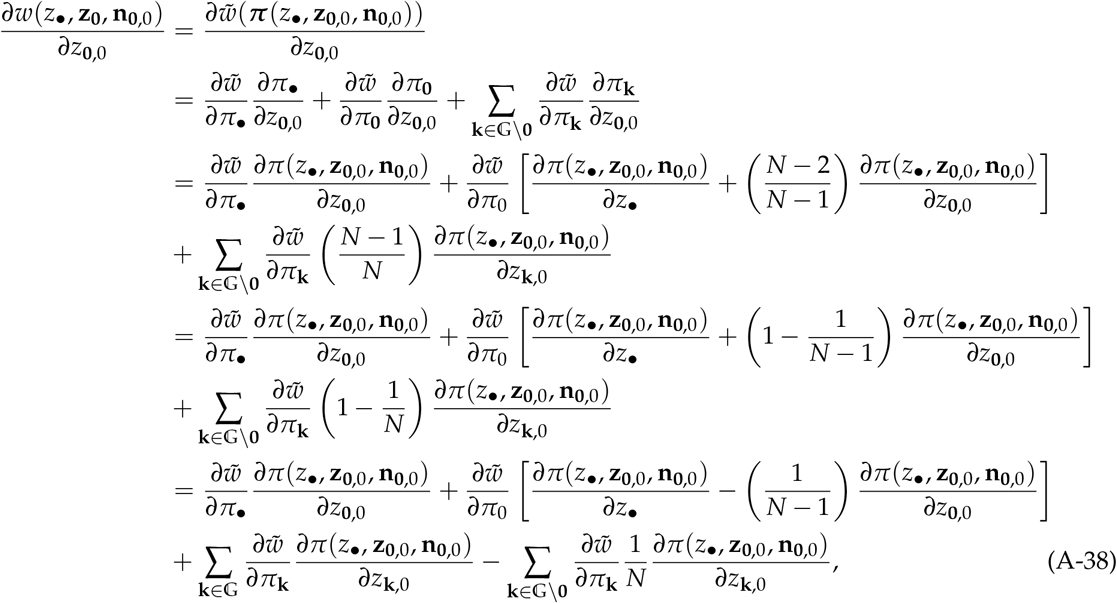

where the first equality follows from taking the derivative to both sides of eq. (A-23); the second equality follows from applying the chain rule; the third equality follows from substituting eqs. (A-28), (A-31) and (A-32); and the last equality follows from distributing and rearranging terms.

Third, the derivative with respect to the average phenotype in any patch **j** ≠ **0** is

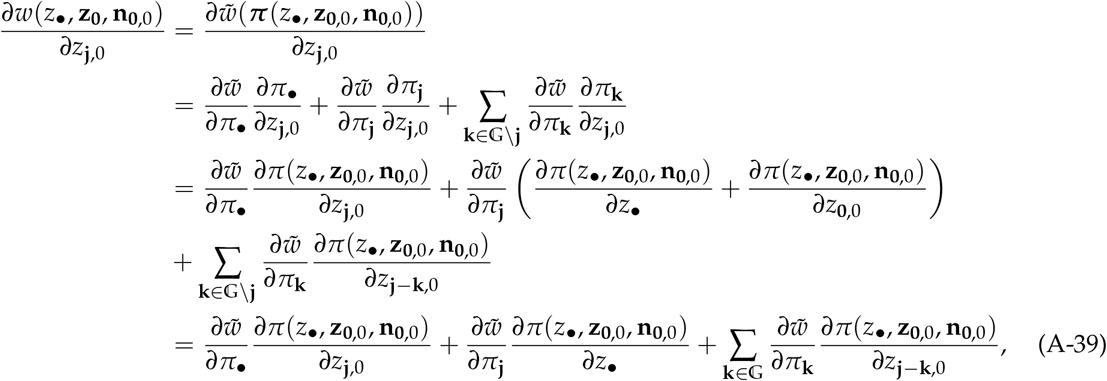

where the second equality follows from applying the chain rule; the third equality follows from using the definition of *π*_•_ (eq. A-24) and substituting eq. (A-33) and eq. (A-34); and the fourth and last equality follows from rearranging.

Finally, the derivative with respect to the state variable in patch **j** is

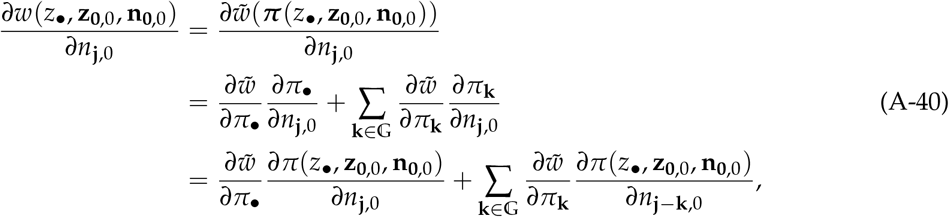

where the second equality follows from applying the chain rule; and the third equality follows from substituting eqs. (A-35)–(A-36).

Let us denote by

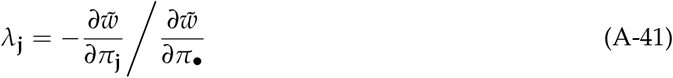

the coefficient of fitness interdependence between individuals in the focal patch and individuals in patch **j**. We can express *s*_w_(*z*) in terms of these coefficients of fitness interdependence and in terms of the derivatives of the fitness function with respect to the phenotypes of different actors in the following way. Substituting eqs. (A-37)–(A-39) into eq. (6a), factoring 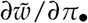, and making use of (A-41), we obtain

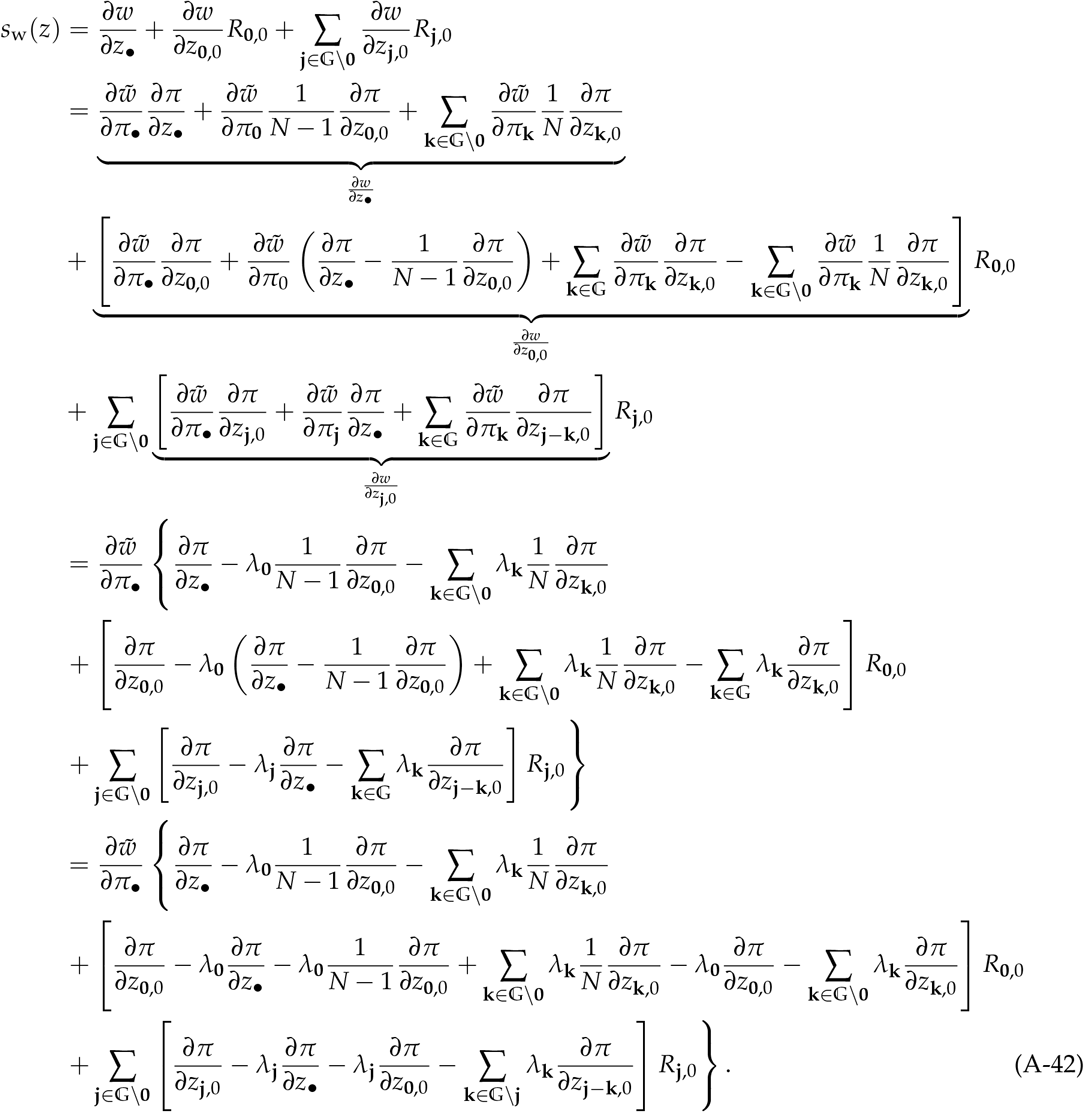

Collecting terms and simplifying, we further get

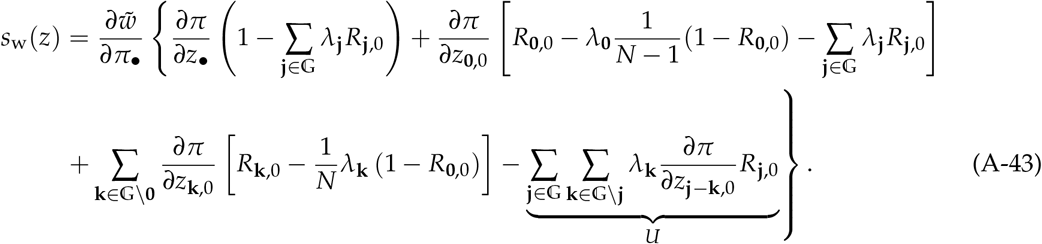

To further simplify this expression, note that the underbraced term can be rewritten as

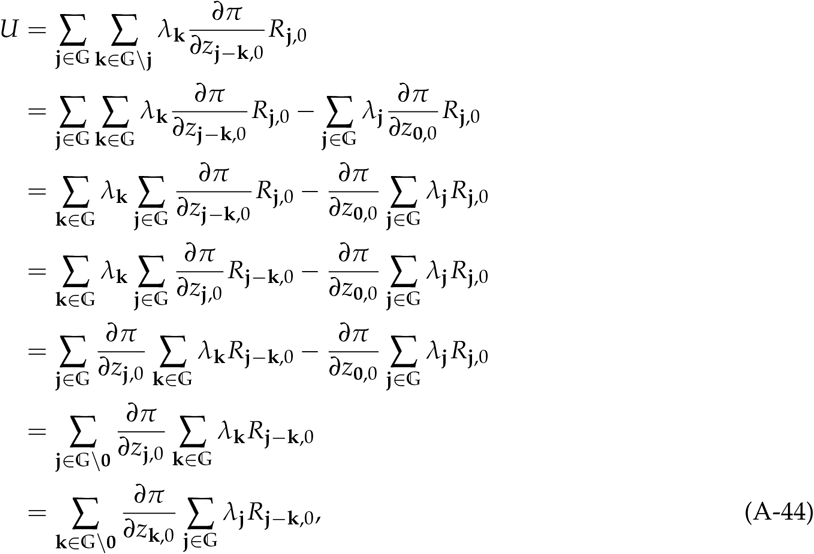

where the third line follows from the identity

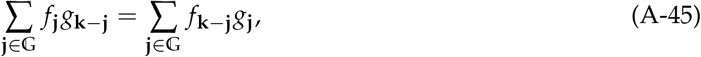

and the last line follows from changing the dummy variables and from the symmetry of the relatedness coefficients (i.e. the fact that *R*_−**k**,0_ = *R*_**k**,0_ holds for all **k** ∈ 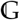).

Substituting (A-44) into (A-43) and simplifying we obtain

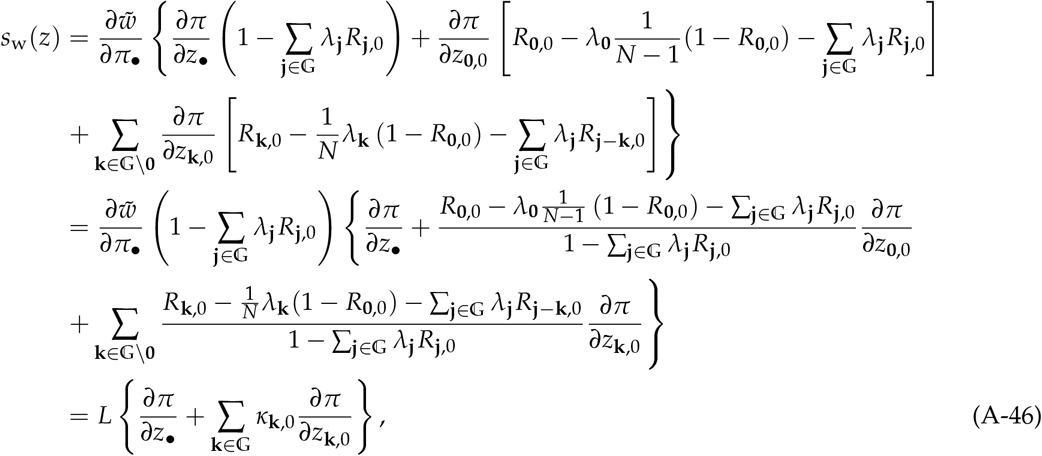

where the second equality follows from factoring 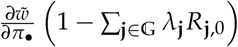, and the final equality follows from defining

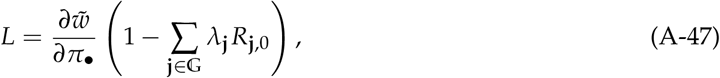

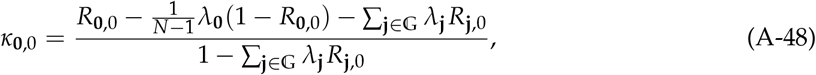

and

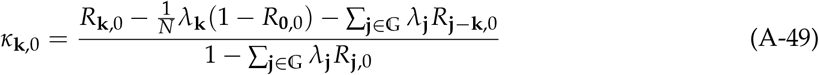

for **k** ≠ **0**. Eq. (A-46) corresponds to the expression for *s*_w_(*z*) in eq. (18) of the main text, as required, by rearranging the numerators of eq. (A-48) and eq. (A-49). The representation of the numerators in eqs. (A-48)–(A-49) are useful for computations, while those in eq. (19) are more amenable to interpretation.

Note that in the infinite island model of dispersal, *R*_**j**,0_ = 0 for all **j** ≠ **0**. In this case, *κ*_**0**,0_ (eq. A-48) reduces to eq. 22 of [7] as it should. This provides a consistency check of our derivation.

Let us turn to express *s*_e_(*z*) in terms of the coefficients of fitness interdependence. From eq. (11) and after substituting (A-40), factoring 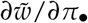 and making use of eq. (A-41), we obtain

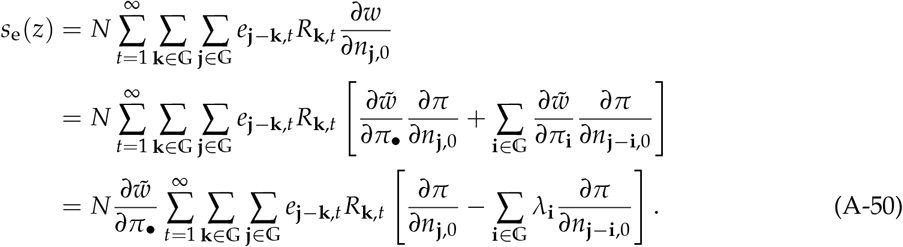

By rearranging terms and applying the identity (A-45), we can rewrite this expression as

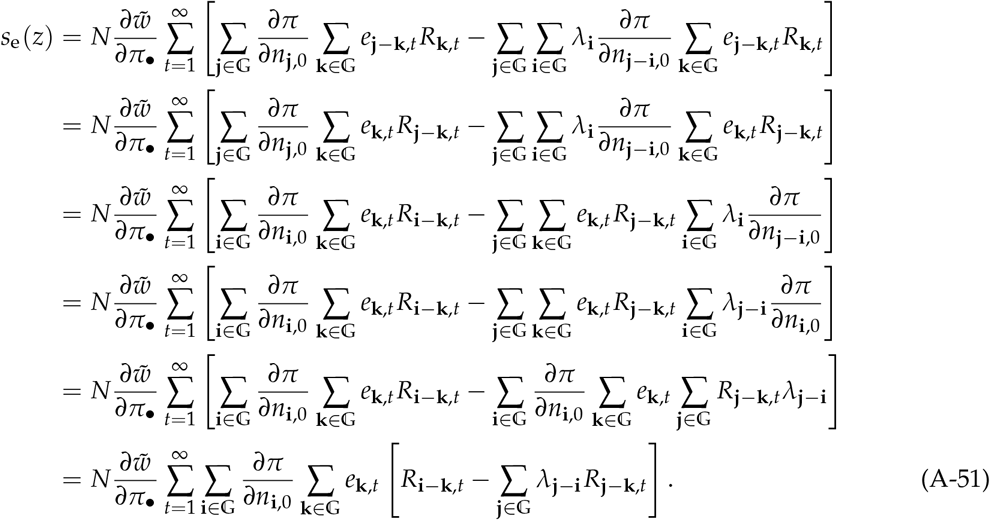

Using the symmetry of the relatedness coefficients and changing the summation indices we further get

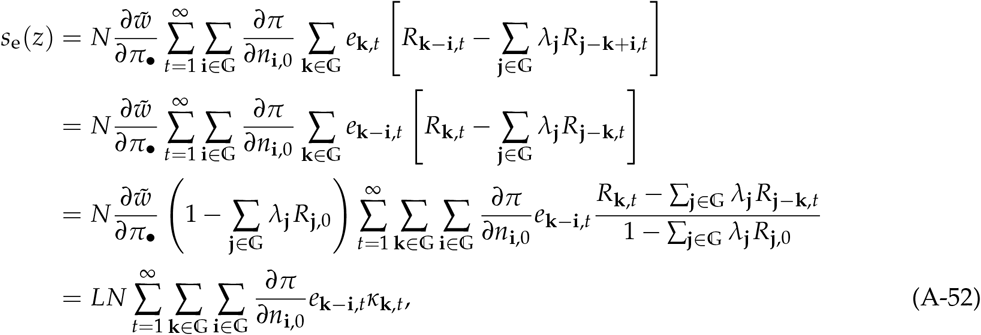

where the second-to-last line follows from multiplying and dividing by 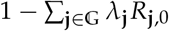, and the last line follows from identifying *L* (A-47) and defining

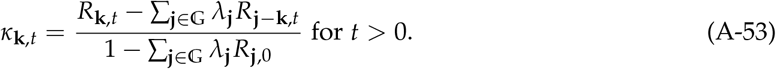

Eq. (A-52) corresponds to the expression for *s*_e_(*z*) in eq. (18) of the main text, as required.

Adding the intra- (eq. A-46) and inter-temporal (eq. A-52) components of the selection gradient, we obtain

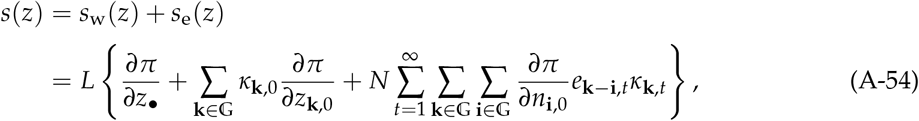

overall.

For derivations to come, it is convenient to have the scaled-relatedness coefficients written in terms of the partial derivatives of 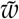 with respect to the payoffs of different individuals. From eqs. (A-48), (A-49) and (A-53) and the definition of the coefficients of fitness interdependence (A-41) we have after rearrangements and multiplying numerators and denominators by 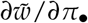:

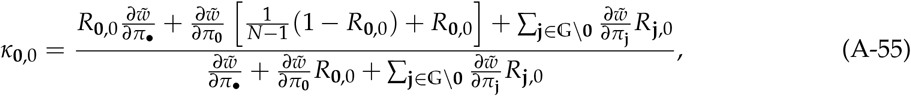

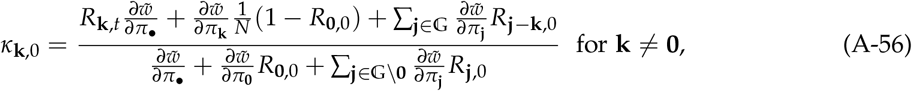

and

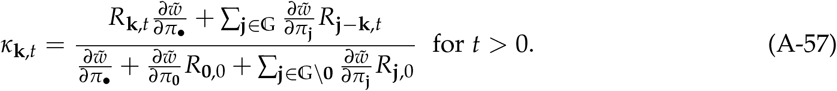

## Appendix E Explicit expressions for scaled relatedness

Here, we derive the explicit expression for scaled relatedness *κ*_**k**,*t*_ given by eq. (III.A) in Box 3, which is based on the assumption that individual fitness can be written as eq. (17); that is

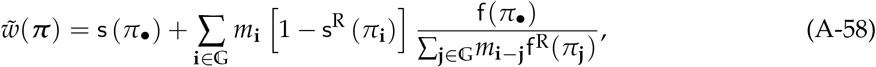

where

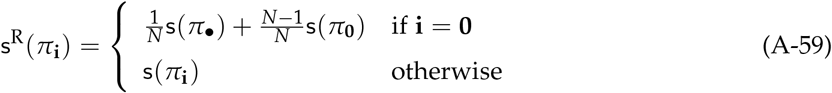

and

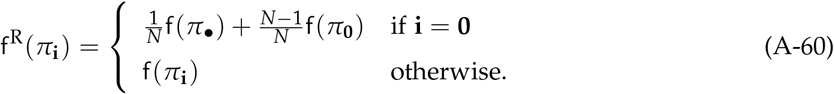

We proceed in three steps. First, we calculate payoff derivatives and the coefficients of fitness interdependence in terms of demographic parameters (Appendix E.1). Second, we calculate expressions for the scaled relatedness coefficients in terms of relatedness coefficients (Appendix E.2). Third, starting from these expressions, we calculate expressions for the scaled relatedness coefficients in terms of demographic parameters, obtaining eq. (III.A) shown in Box 3 (Appendix E.3). Finally, in Appendix E.4, we use these results to get an expression for *L* (A-47) in terms of demographic parameters, which can be useful to have the magnitude (not just the sign) of the selection gradient.

### Appendix E.1 Payoff derivatives and coefficients of fitness interdependence

Using the quotient rule of derivatives, and evaluating expressions at the resident trait value, the derivative of 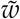(A-58) with respect to the payoff of the focal individual *π*_•_ can be written as

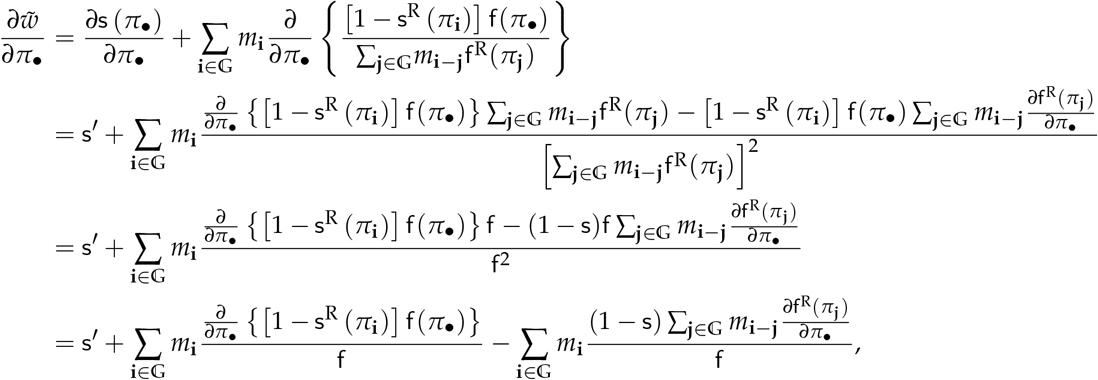

where we have set s^′^ = *∂*s (*π*_•_) /*∂π*_•_, and used the fact that 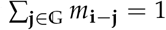 for all **i** ∈ 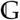. Substituting (A-59), noting that

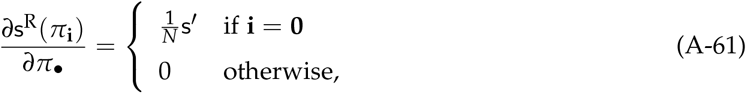

and

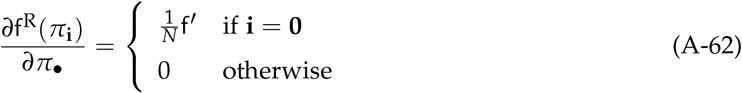

hold, and setting f^′^ = *∂*f (*π*_•_) /*∂π*_•_, we further get

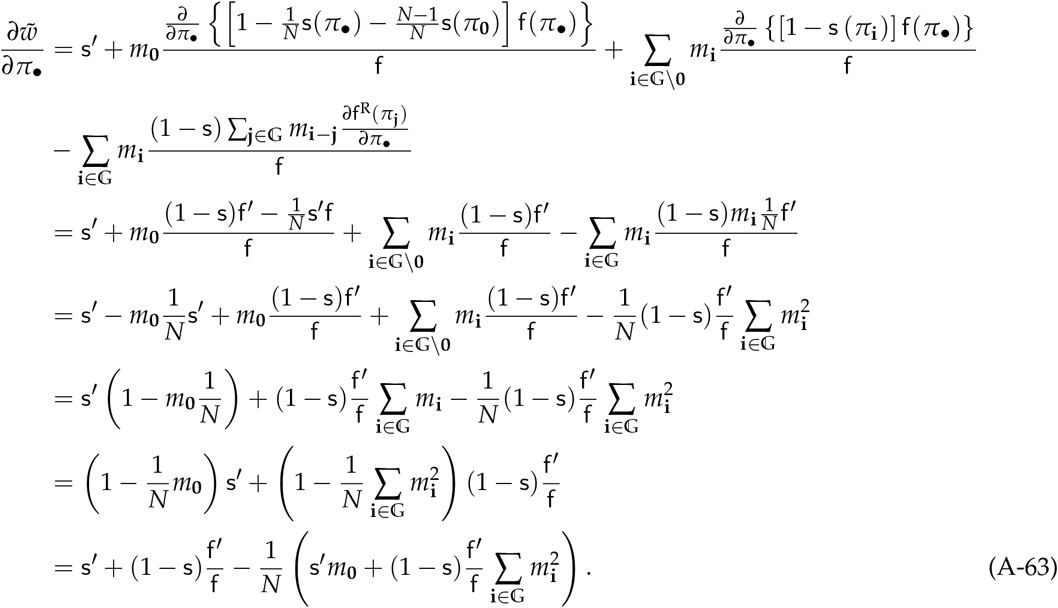

Applying the same line of arguments produces

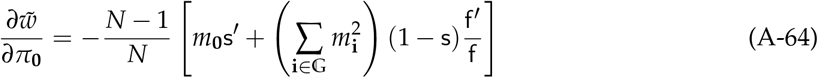

and

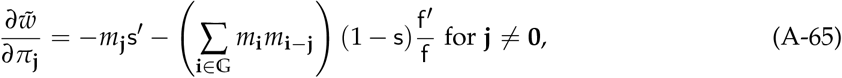

where, as usual, all functions are evaluated at the resident trait value *z* and equilibrium 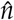.

Introducing the notation

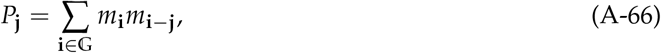

which is the probability that an offspring born in patch **j** competes with an offspring of the focal individual (i.e. that they both migrate to the same patch), the derivatives in eqs. (A-63)–(A-65) can be more compactly written as

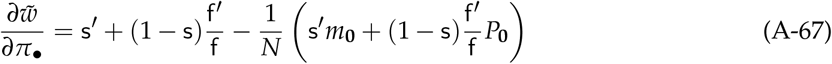

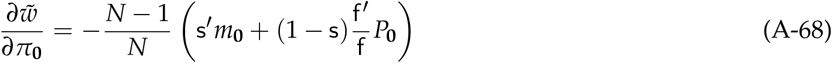

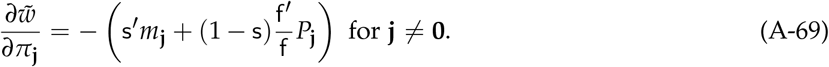

In terms of these derivatives, the coefficients of fitness interdependence (A-41) can then be written as

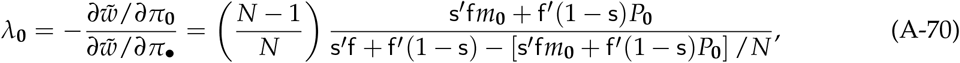

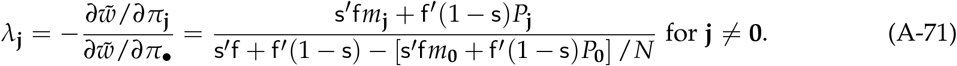

### Appendix E.2 Scaled-relatedness in terms of relatedness coefficients

To calculate and simplify the scaled-relatedness coefficients, it is convenient to start from expressions (A-55) – (A-57). First, note that using eqs. (A-67) and (A-68), and rearranging terms yields

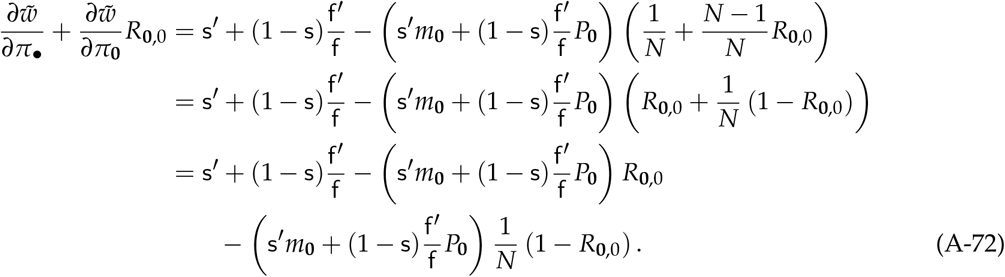

Then, using (A-72) and (A-69), rearranging, and factoring, the common denominator of *κ*_**0**,0_ (A-55), *κ*_**k**,0_ (A-56), and *κ*_**k**,*t*_ (A-57) can be written as

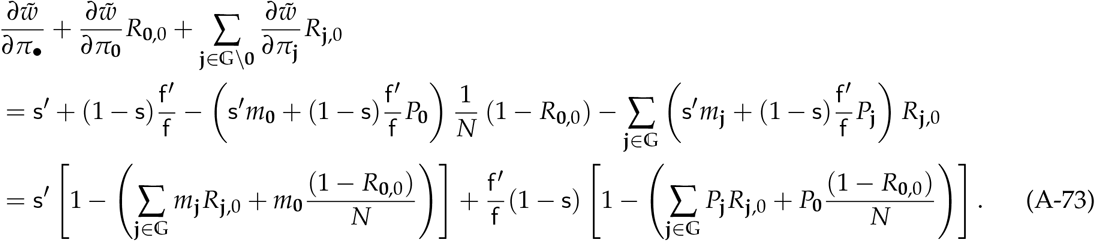

Similarly, using (A-68), yields

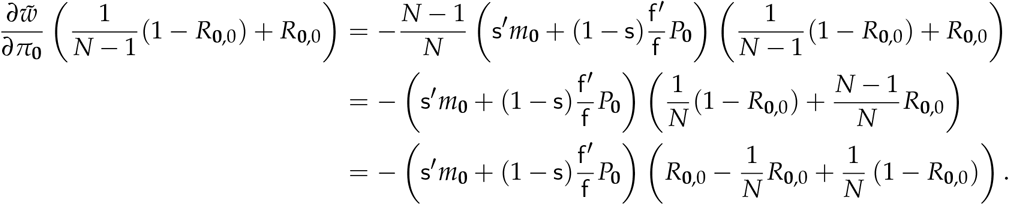

Using this expression together with (A-67) and (A-69), the numerator of *κ*_**0**,0_ (A-55) can be simplified as follows:

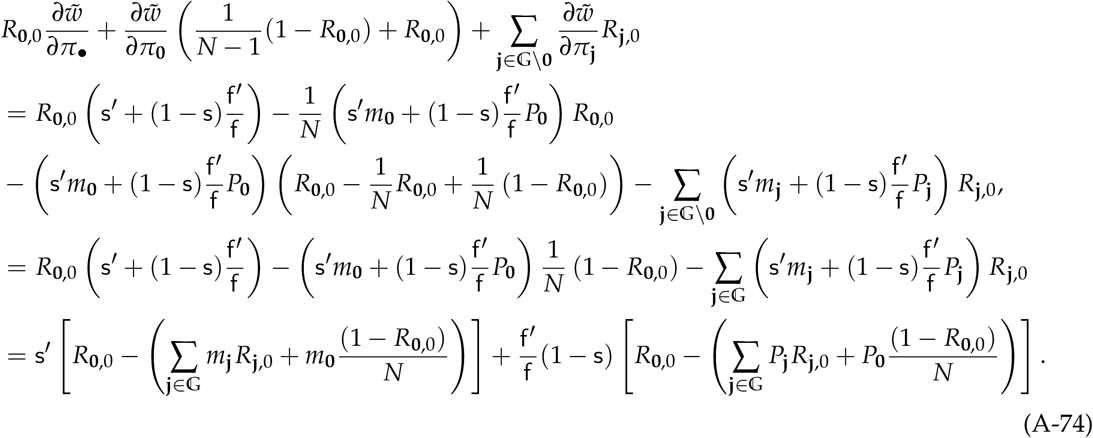

Now, substituting (A-73) and (A-74) into (A-55), and then multiplying numerator and denominator by f, yields

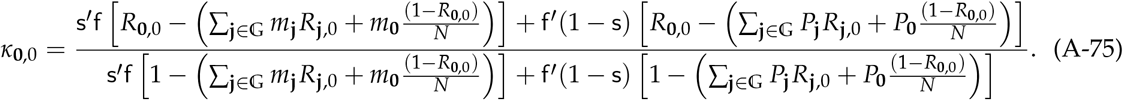

To obtain similar expressions for *κ*_**k**,0_ (A-56), and *κ*_**k**,*t*_ (A-57), we first use eqs. (A-67)–(A-69) to write

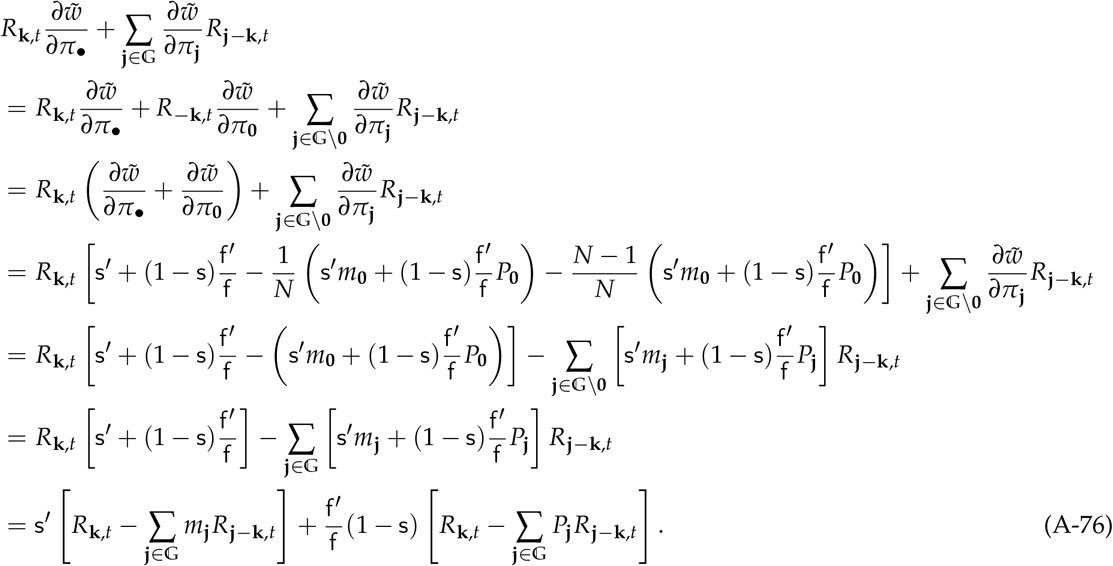

and

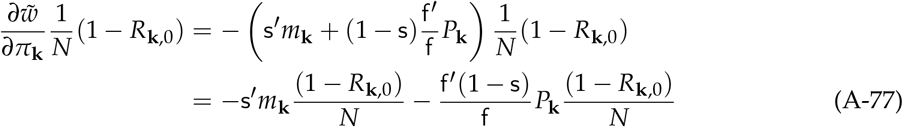

Substituting (A-73), (A-77), and (A-76) into (A-56), and then multiplying numerator and denominator by f, yields

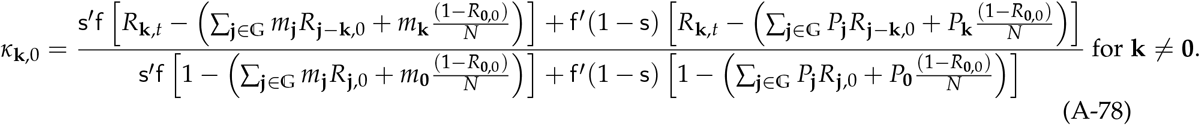

Likewise, substituting (A-73), and (A-76) into (A-57), and then multiplying numerator and denominator by f, yields

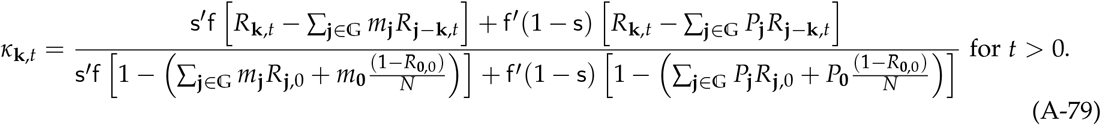

### Appendix E.3 Scaled-relatedness in terms of demographic parameters

Substituting eq. (7) into eqs. (A-75), (A-78), and (A-79), and then cancelling common terms, we obtain

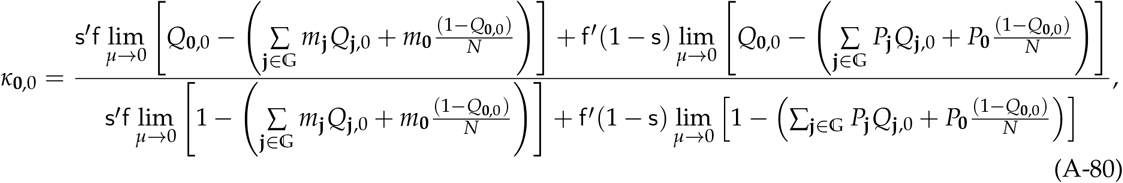

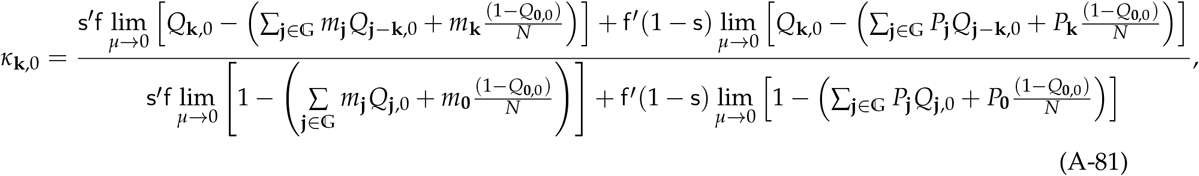

for **k** ≠ **0**, and

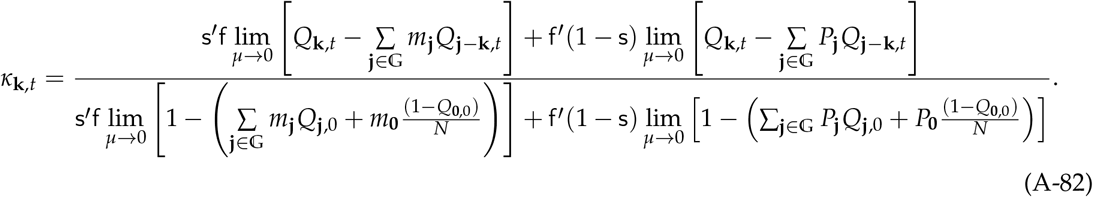

for *t* > 0.

To simplify eqs. (A-80)–(A-82), we first note that, from eqs. (A.42), (A.48), and (A.51) in [8], we have^1^

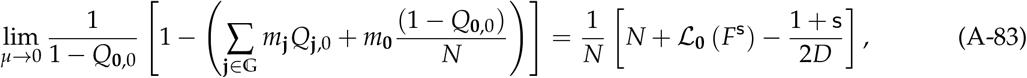

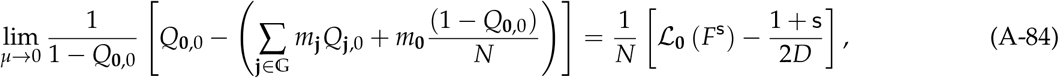

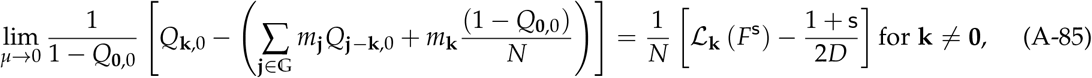

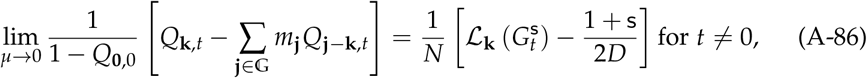

where ℒ_**k**_(ℱ) is the inverse transform of ℱ at **k** as defined in eq. (I.B), and where the functions *F*^s^ and 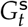 are defined at **h** as

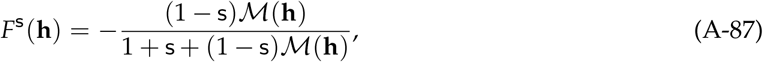

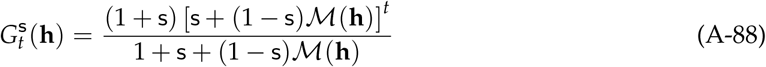

(see eqs. (A.47) and (A.49) in [8]).

Likewise, from eqs. (A.32), (A.38), and (A.41)^2^ in [8], we have

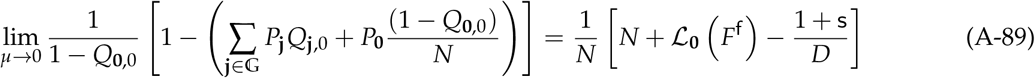

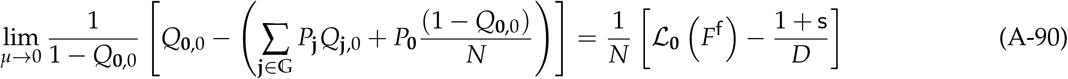

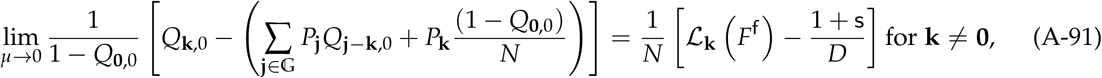

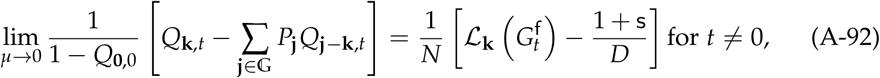

where the functions *F*^f^ and 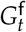 are defined at **h** as

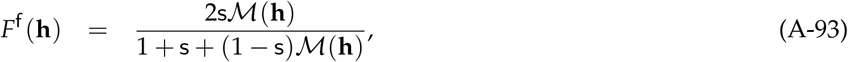

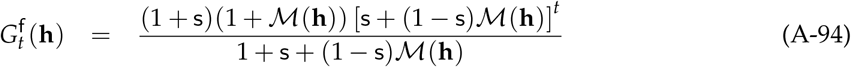

(see eqs. (A.37) and (A.39)^3^ in [8]).

We can now proceed to simplify eqs. (A-80)–(A-82). First, multiplying the numerator and denominator of (A-80) by lim_*μ*→0_ 1/(1 − *Q*_**0**,0_), substituting eqs. (A-83), (A-84), (A-89), and (A-90), and then multiplying numerator and denominator by *N*, we obtain

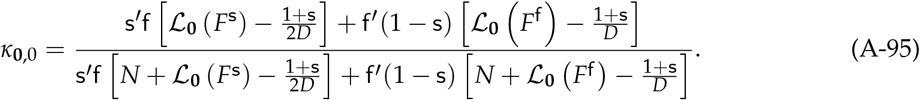

Second, proceeding similarly with eq. (A-81) (by substituting eqs. (A-83), (A-85), (A-89), and (A-91)), we obtain

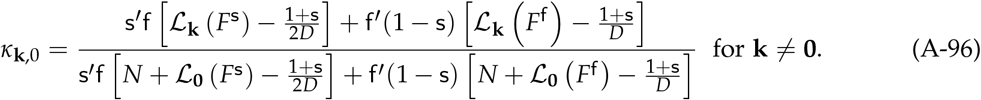

Third, proceeding similarly with eq. (A-82) with *t* > 0 (by substituting eqs. (A-83), (A-86), (A-89), and (A-92)) we obtain

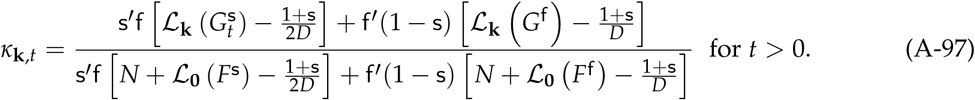

Finally, noting that eq. (A-95) is equal to eq. (A-96) with **k** = **0**, substituting eqs. (A-87)–(A-93), and rearranging yields eq. (III.A) of Box 2, that is:

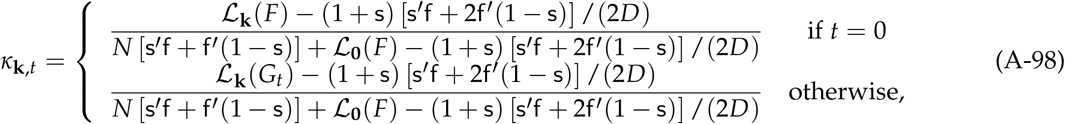

where the functions *F* and *G*_*t*_ are given by

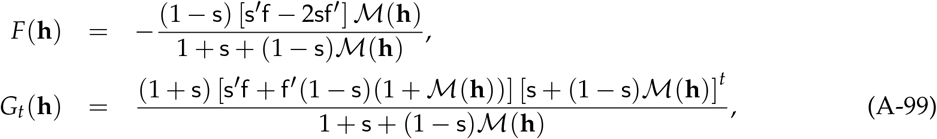

as required.

Note that one can also write eq. (A-98) as

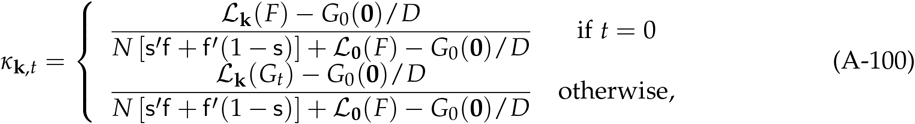

since, for all *t*,

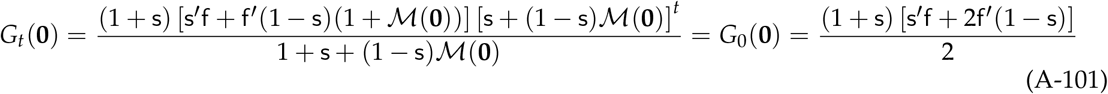

holds.

### Appendix E.4 Explicit expression for *L*

Finally, we evaluate *L* in eq. (A-47), which is needed if one aims to evaluate the trait stationary density function (A-5). Substituting the definition of the coefficients of fitness interdependence (A-41) into eq. (A-47), simplifying, and rearranging, we obtain

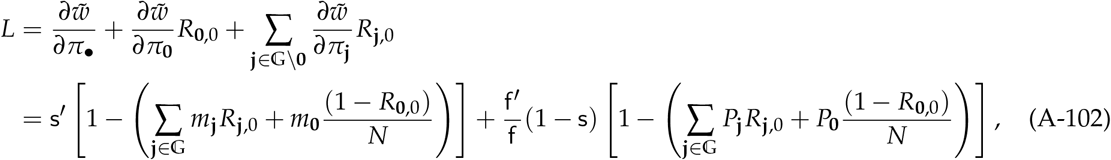

where the second equality follows from our previous derivation in eq. (A-73).

Substituting eq. (7), we get

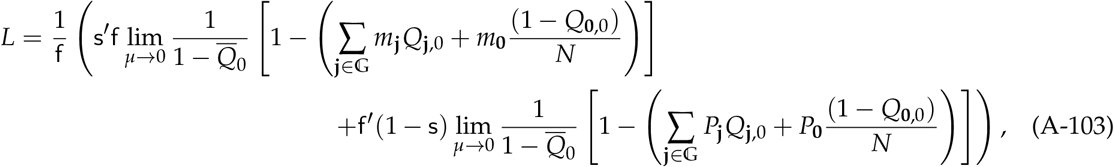

which can be computed as

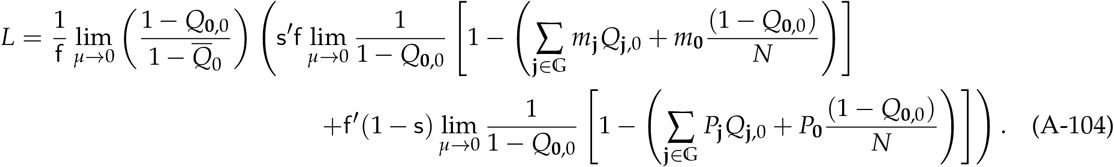

Using eqs. (A-83), (A-89), and (A-99), this becomes after some rearrangements,

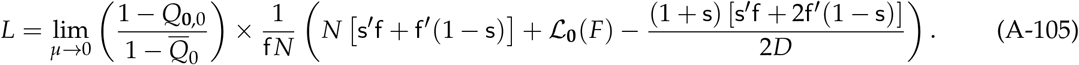

For a Wright-Fisher process where s^′^ = s = 0 (and hence, also ℒ_**0**_(*F*) = 0), we have

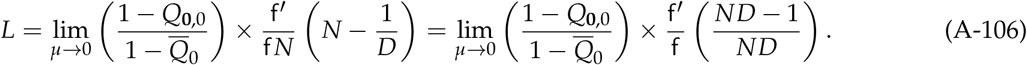

## Appendix F Explicit coefficient for the selection gradient

Here, we derive eq. (III.C) of Box 3 of the main text. First, we simplify the expression for *κ*_**k**,*t*_ for *t* > 0 given in the second line of eq. (A-100). Using the definition of the inverse Fourier transform given in eq. (I.D), and simplifying we obtain

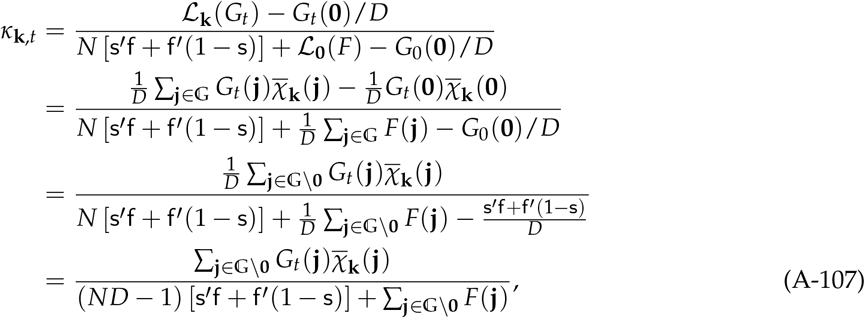

where we have used 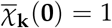 for all **k** ∈ 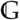, the identity *G*_*t*_(**0**) = *G*_0_(**0**) for all *t* (A-101), and the fact that

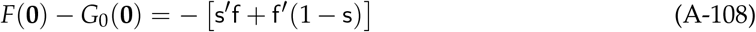

holds.

Substituting the simplified expression for *κ*_**k**,*t*_ (A-107) together with the expression for *e*_**k**,*t*_ (14) into eq. (24), and setting

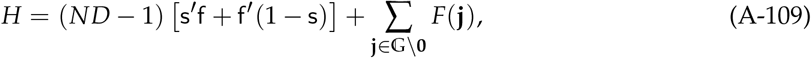

Yields

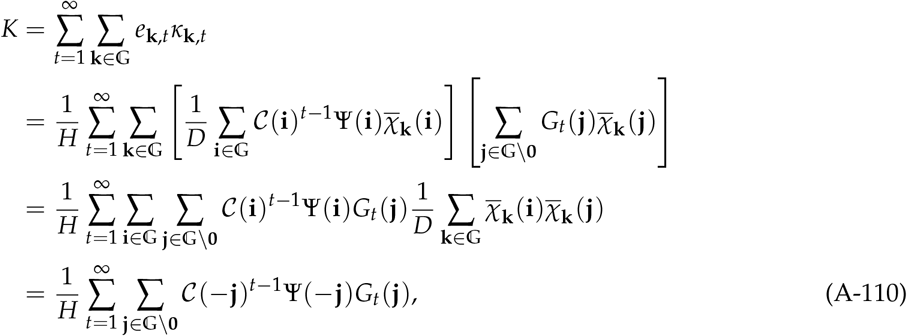

where the last equality follows from using eq. (I.F). Substituting eq. (III.B) into (A-110) and solving the geometric series yields

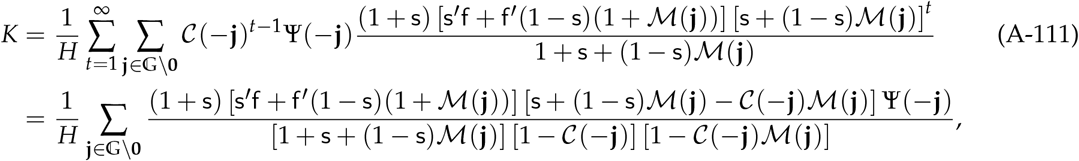

which is the final expression presented in eq. (III.C).

To go from the first to the second line of eq. (A-111), the relevant geometric series must converge, which happens if the moduli of ℳ(**j**) and 𝒞(**j**) are smaller than one (i.e. |ℳ(**j**)| < 1 and |𝒞(**j**)| < 1) for all **j** ≠ **0**, i.e. if the complex numbers ℳ(**j**) and 𝒞(**j**) are within the unit circle. To see this is true, consider first that by the property of characteristic functions of probability distributions, we have ℳ(**0**) = 1, and |ℳ(**j**)| < 1 for **j** ≠ **0** (p. 182 in [9]). Second, |𝒞(**j**)| < 1 from our assumption that the dynamical system eq. (2) has a hyperbolically stable equilibrium point. Indeed, stability means that all the eigenvalues of the Jacobian matrix of eq. (2) have modulus smaller than one (e.g. p. 103 of [10]). But these eigenvalues are in fact given by the coefficients C(**j**). To see this, first note that from eq. (2) the Jacobian of this discrete-time dynamical system around the equilibrium 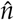 defined by eq. (3) is given by

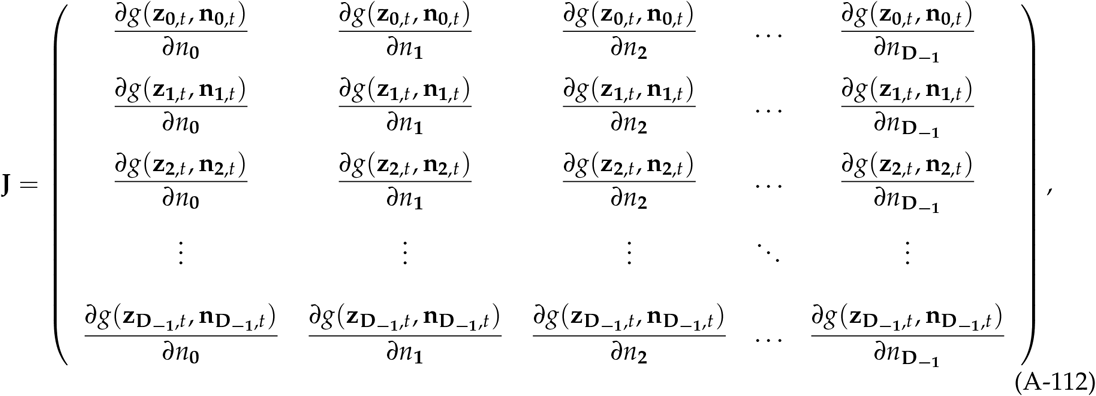

where all derivatives are evaluated at *z* and 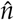. Now, recalling the notations defined in eq. (A-20), the entries of this matrix are of the form

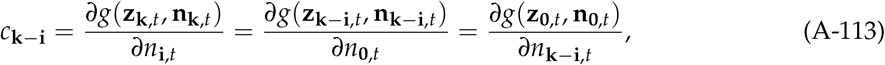

which is the same as eq. (A-20) since all phenotypes vectors, here and there, are set to (*z*, …, *z*) when computing the derivative. From the first equality in the previous equation, the Jacobian (A-112) can be written as

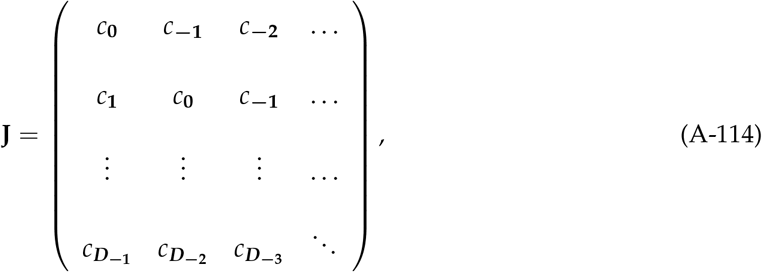

where we defined ***D***_−2_ = **D**_−1_ − **1, *D***_−3_ = **D**_−1_ − **2**, etc. Written in this form, it is clear that the Jacobian (A-114) is a 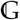 -group circulant matrix (e.g. p. 50 of [11]), with eigenvalues given by the Fourier transform of *c*_**j**_ (Theorem 8 in [11]). Hence, the **k**-th eigenvalue of **J** is 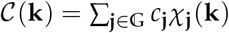.

## Appendix G Public good diffusion example

### Appendix G.1 Fecundity effects

Here, we derive eq. (33) of the main text, which considers fecundity effects and no generational overlap (s^′^ = s = 0). Substituting eq. (21) into eq. (31) yields

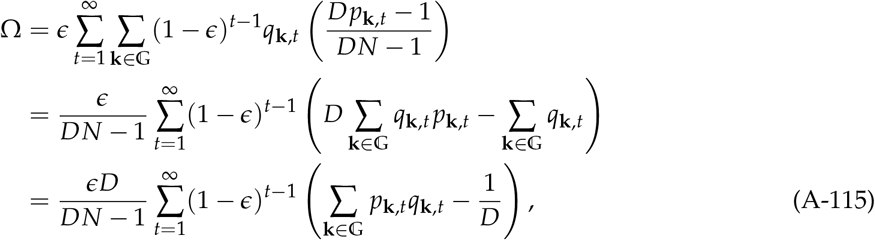

where we have used the fact that *q*_**k**,*t*_ is a probability distribution over 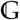 for all *t* and hence that 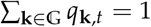 holds for all *t*.

Eq. (A-115) can be written in terms of the population covariance of *p*_**k**,*t*_ and *q*_**k**,*t*_ in the following way. Recall that the population covariance of two vectors *x* = (*x*_1_, …, *x*_*n*_) and *y* = (*y*_1_, …, *y*_*n*_) of length *n* is given by

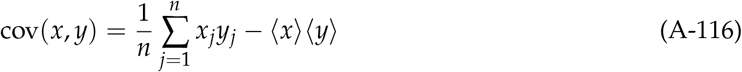

where 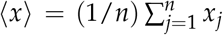 and 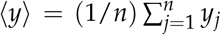. Using this definition of population covariance, and denoting by 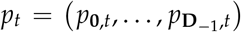 and 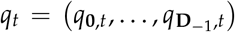 the vectors collecting all *p*_**k**,*t*_’s and *q*_**k**,*t*_’s in lexicographic order, we can write

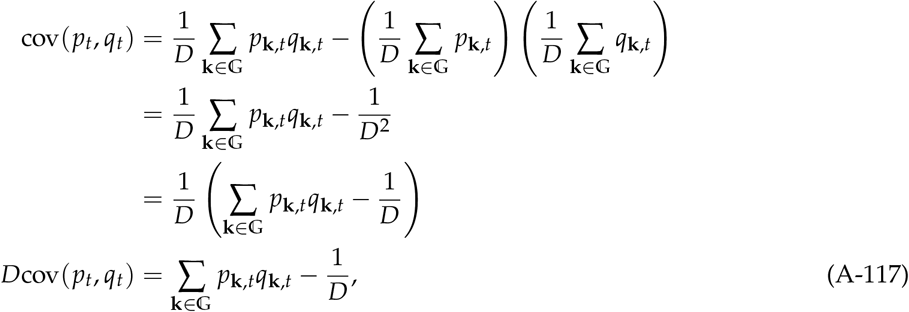

where the second line follows from the fact that both *p*_**k**,*t*_ and *q*_**k**,*t*_ are probability distributions over 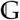 for all *t* and hence satisfy 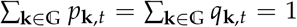 for all *t*. Substituting (A-117) into (A-115) we finally obtain

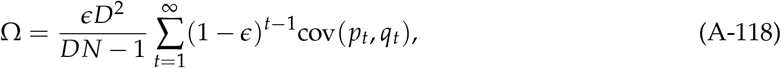

as required.

### Appendix G.2 Fecundity effects: Weak dispersal

Here, we derive eq. (35) of the main text, following the common approach to evaluate a weak migration approximation (chapter 3 in [2]). To do so, we first set *m*_**0**_ = (1 − *m*) and *d*_**0**_ = (1 − *d*), where *m* and *d* are the net dispersal probabilities of the focal species and the environmental variable, and write 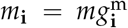 and 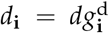. The characteristic functions of the dispersal distributions can then be expressed as ℳ(**j**) = 1 − *mx*^m^(**j**) and 𝒟(**j**) = 1 − *dx*^d^(**j**), where 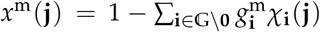 and 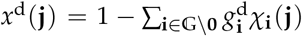. Substituting these expressions into the summand of eq. (34), and Taylor expanding around *m* = 0 and *d* = 0, we get

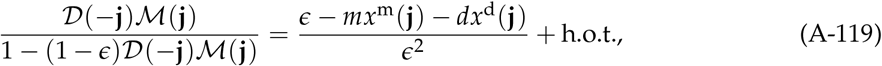

where “h.o.t.” refers to higher order terms, e.g. terms proportional to *m*^2^, *md, d*^2^, etc. Substituting *x*^m^(**j**) = [1 − ℳ (**j**)]/*m* and *x*^d^(**j**) = [1 − 𝒟 (**j**)]/*d*, we can write eq. (34) as

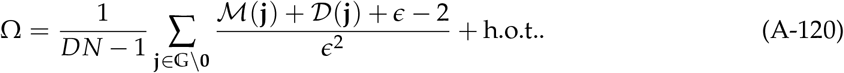

Neglecting the higher order terms and using 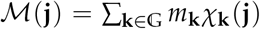 and 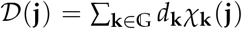 produces

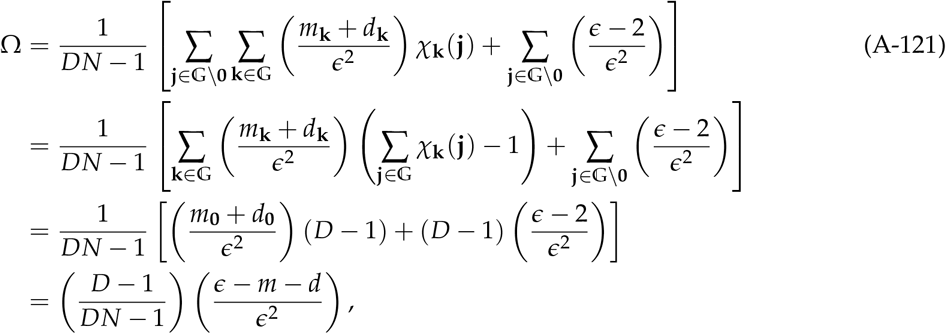

where the penultimate equality follows from the facts that 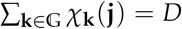 if **j** = **0** and zero otherwise (recall eq. I.F), and that 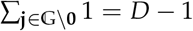.

### Appendix G.3 Fecundity effects: Island model

Here, we derive the expression for Ω for a Wright-Fisher life cycle (using eq. 34) under the island model of species dispersal and commons movement, which we use in Fig 6 (dashed lines). Under this model of dispersal and movement, we have

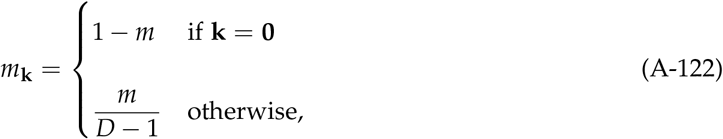

and similarly,

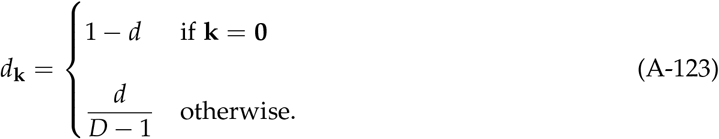

We first calculate the Fourier transforms ℳ(**h**) and 𝒟( − **h**) of the dispersal and movement distributions. We obtain

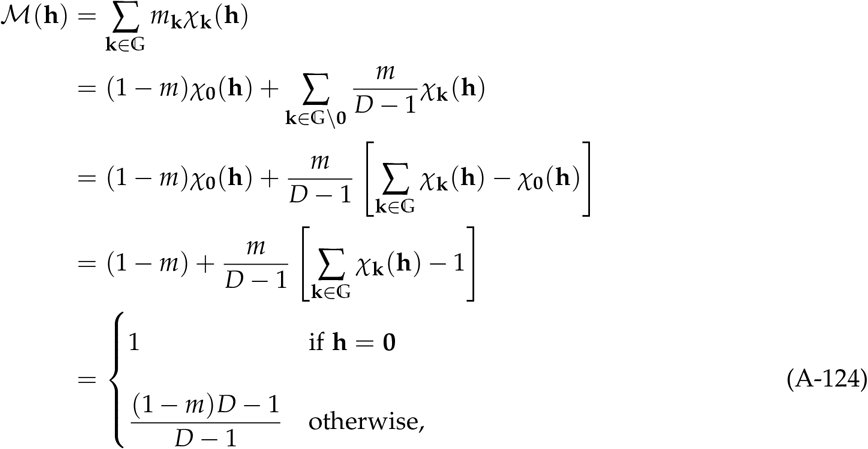

where the first line uses the definition of Fourier transform (I.B), the second and third lines rearrange terms, the fourth line uses the fact that *χ*_**0**_(**h**) = 0, and the fifth line follows from identity (I.F). Likewise,

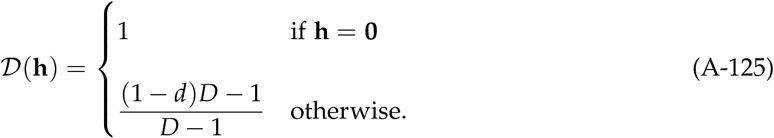

Substituting eqs. (A-124) and (A-125) into eq. (34), using the fact that, for (A-125) 𝒟 ( − **h**) = 𝒟 (**h**) holds, and simplifying, we arrive to

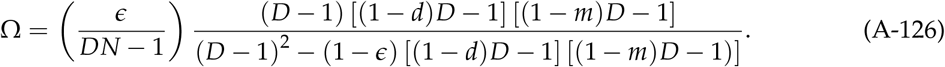

### Appendix G.4 Survival effects

We now consider the case where there are survival but no fecundity effects (f^′^ = 0). Writing Ω as Ω = *ϵKN*/*P*^′^(*z*), substituting

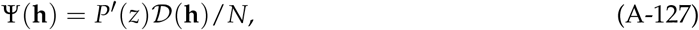

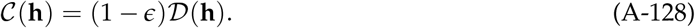

into eq. (III.F) in Box 3, and simplifying we obtain

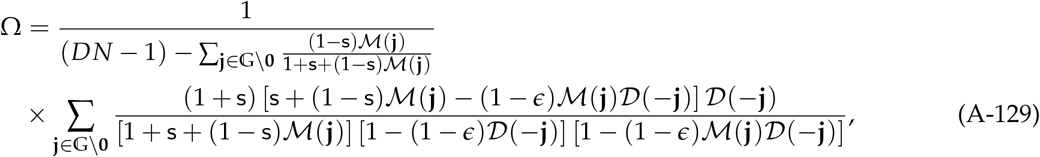

which remains a somewhat complicated expression. In the limit s → 1, eq. (A-129) simplifies to

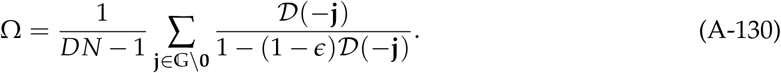

This can be thought of as a special case where investment into the common-pool resource occurs in a population of immortal individuals that, therefore, become mortal through endogenously induced deaths. Finally, we note that for a spatially symmetric dispersal distribution we can set 𝒟( − **j**) = 𝒟(**j**).

### Appendix G.5 Species dispersal and commons movement

In our example, we assumed that the evolving species dispersed according to a model based on the binomial distribution, which is detailed in Appendix B. Hence, the characteristic function used for a one-dimensional habitat is given by eq. (A-10), while for a two-dimensional habitat, it is based on Appendix B.2.

We assume that the way the commons moves in space follows the same model as the evolving species. We write *d* for the commons’ probability of movement (instead of *m*), and 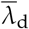 for the mean number of steps a unit of commons moves conditional on leaving the patch (instead of 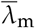). The characteristic function of the movement in one dimension then is like eq. (A-10), i.e.

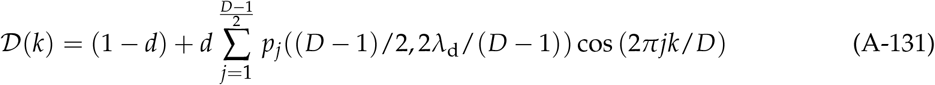

where *λ*_d_ is such that

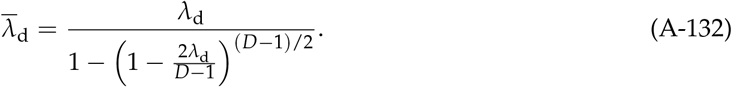

### Appendix G.6 Stationary distribution

Here, we specify the stationary distribution of the trait substitution sequence for our example, i.e. we specify eq. (A-5), which we used in Fig 6B for the interval. Substituting eq. (30) into eq. (A-54), which is in turn substituted into eq. (A-4) gives

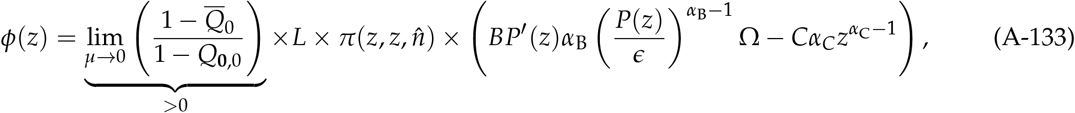

thus characterising the term within parenthesis of eq. (A-5). For the Wright-Fisher process, we have from eq. (A-106) that

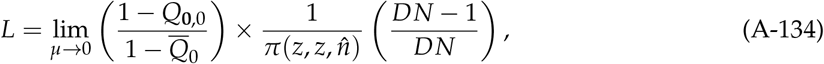

where we used the fact that, for our example, payoff is fecundity, and so f^′^ = 1 in eq. (A-106). Thus, the perturbation of the fixation probability reduces to

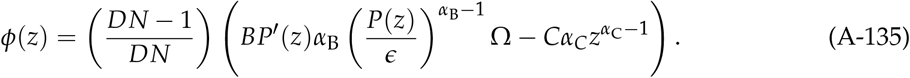

Assuming further that *P*(*z*) = *P*_0_*z*, we find by substituting eq. (A-135) into eq. (A-5) that the stationary distribution is given by

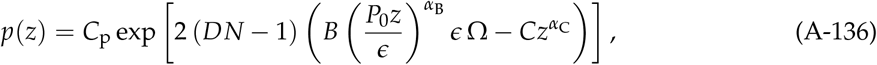

where *C*_p_ is a constant of proportionality such that 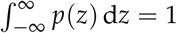.

## Footnotes

1 eq. (A.51) in [8] applies for all **k** ∈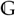, the condition “if **k** > **0**” therein is not necessary. Also, the term − *s*/*N* in the last line of eq (A.48) of [8] contains a typo and should be replaced by *s*/*N*.

2 eq. (A.41) in [8] applies for for all **k** ∈ 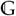, the condition “if **k** ≠ **0**” therein is not necessary.

3 eq. (A.39) in [8] contains a typo in that the the second parenthesis is not closed and the term (1 + *ψ*_**h**_ should read (1 + *ψ*_**h**_).

